# Perinatal Lead (Pb) Exposure Increases Mouse Embryonic Weight and Alters Neuronal Gene Expression

**DOI:** 10.1101/2025.09.18.677210

**Authors:** Bambarendage P. U. Perera, Minghua Li, Anagha Tapaswi, Junru Pan, Dongyue Wang, Tejas Goswami, Rachel K. Morgan, Kelly M. Bakulski, Jaclyn M. Goodrich, Maureen A. Sartor, Dana C. Dolinoy, Justin A. Colacino

## Abstract

Acute and chronic exposure to lead (Pb) during pregnancy is linked to adverse health outcomes, including delayed neurodevelopment in offspring. However, the pathways by which Pb exposure influences long-term health remain poorly understood. To address this, we measured the effects of perinatal Pb exposure on gene expression including imprinted genes, X-linked genes, and sexually dimorphic genes. Female mice were given control or Pb acetate dosed (32 ppm) drinking water two weeks prior to timed mating until embryonic day (E)10–12, upon which whole embryos were collected, weighed, and sexed at E13–15. From a subset of embryo heads (*n*≥9 per sex per group), we extracted and sequenced RNA. We used linear regression to assess Pb impacts on embryonic weight and gene expression across all mice and stratified by sex. Among the differentially expressed genes, we identified significantly enriched pathways. Pb-exposed embryos weighed more than controls (*p*=0.007), across both sexes. Collectively, we identified 2,920 differentially expressed genes (FDR<0.05), including 31 imprinted genes and 120 X-linked genes upon Pb exposure. Pb exposure altered expression in gene pathways related to neuronal structure and function as well as sexually dimorphic genes (44 for females; 76 for males). These findings highlight perinatal Pb-linked alterations that may drive later-life health outcomes.

## 1. INTRODUCTION

Environmental and occupational lead (Pb) exposure impacts human populations in the US and around the world and has neurotoxic, cardiovascular, and metabolic effects ^1,2^. Sources of Pb exposure include leaded paint, pipes, electronics, industrial activities, and contaminated soil and water. There is no known ‘safe’ level of Pb exposure. An estimated 500,000 children under the age of 6 in the U.S. are reported to have ≥5 μg/dL concentrations of blood Pb ^3^. Children and pregnant women are particularly susceptible to the health effects of Pb, with gestational exposure linked to adverse birth outcomes, delayed neurodevelopment, growth defects, and potential contributions to long-term neurological health issues such as Alzheimer’s disease ^4–6^. Despite abundant literature on Pb-induced toxicity, the gene pathways driving its health effects, particularly during early stages of neuronal development, remain poorly understood.

According to the Developmental Origins of Health and Disease hypothesis, environmental exposure during development predisposes offspring to diseases later in life ^7^. Early development is largely driven by the epigenome, which regulates gene expression via DNA methylation (DNAm), histone modifications, and non-coding RNA (ncRNA), and is highly sensitive to environmental exposures. In therian mammals, including placental mammals and marsupials, offspring inherit one maternal allele and one paternal allele, giving rise to biallelic autosomal gene expression. However, a subset of these genes is exclusively expressed from one parental allele, resulting in parent-of-origin specific monoallelic expression through the epigenetically regulated process of genomic imprinting ^8^. Dysfunctional genomic imprinting is implicated in several human diseases such as Angelman syndrome and Prader-Willi syndrome ^9–12^. For instance, loss of function of the maternally-inherited ubiquitin protein ligase E3A (*UBE3A*) contributes to the onset of Angelman syndrome, while the loss of paternally expressed genes including small nuclear ribonucleoprotein polypeptide N (*SNRPN*) contributes to the onset of Prader-Willi syndrome ^13–15^. Imprinted genes are highly sensitive to the environment during early gestation and are critical for proper embryonic growth and development ^16–19^. Human cohort studies and perinatal Pb exposure mouse models support that Pb exposure can alter imprinted genes and related mechanisms ^20–22^.

Several reported effects of Pb neurotoxicity are sex-specific, with notable changes in gene expression and epigenetically regulated mechanisms ^23–26^. Among the latter is the regulation of X-linked genes, which are located on the X chromosome and have differing copy numbers between females and males ^27^. Therian mammals undergo X-chromosome inactivation (XCI) as dosage compensation for balancing the X-linked genes in females ^28^. Therefore, X-linked genes are critical for proper brain development and neuronal functions, with XCI being established during the early stages of embryogenesis and extensively studied in mouse models ^27^. XCI is a complex epigenetic mechanism involving the expression of the X-inactive specific transcript (*XIST*) long ncRNA (lncRNA) among others to facilitate imprinted and random XCI in females ^28^. The process of XCI may be modulated by environmental cues, potentially leading to skewed XCI and altered gene expression. However, the impact of Pb exposure on X-linked genes is not fully understood. Developmental exposure to Pb has been linked to sex-dependent effects in human and animal studies ^29,30^. Sexual dimorphism refers to the species-specific differences between females and males in their physical or biological characteristics ^31^. Pb alters DNAm signatures in a sex-specific manner *in vivo*, impacting autosomal genes linked to neurodevelopment, cardiovascular disease, and immune functions differently in sexes ^23,30,32^. These molecular changes may be modulated by sex hormones, contributing to differential disease susceptibility and phenotypic outcomes in brain and behavior between the sexes following Pb exposure ^33^.

We hypothesized that Pb exposure impacts gene expression including of imprinted genes, X-linked genes, and sexually dimorphic autosomal genes. The current work evaluated offspring brain-specific transcriptomic changes associated with Pb exposure using a mouse model of prenatal and gestational exposure using embryonic heads at E13–15. The study identified differentially expressed genes, differentially expressed imprinted genes, differentially expressed X-linked genes, sexually dimorphic genes, and enriched pathways among these genes associated with Pb exposure. The current study is the first to report that Pb exposure during preconception and early gestation is associated with increased mid-gestation embryonic weight, and that sexually dimorphic genes are particularly sensitive to Pb exposure in the mouse nervous system. This work will lay the foundation for developing targeted interventions to alleviate long-term health consequences of early life exposure to Pb.

## 2. MATERIALS AND METHODS

### 2.1. Animal exposure study design

C57BL/6J adult female mice at 10–12 weeks of age were randomly selected and exposed to either control or Pb acetate drinking water (pH 4.5; adjusted using glacial acetic acid) at 32 ppm starting 2 weeks before mating (preconception) and continued until E10–12, and euthanized at E13–15 ^26,34^. All animals were maintained on a phytoestrogen-free modified AIN-93 G diet (Td.95092, 7% corn oil diet, Envigo) while housed in polycarbonate-free cages with 12-hour light and dark cycles ^25,26,34^. Study dams were time-mated by assembling randomly selected 3-month-old males in breeding cages for approximately 48 hours (spanning three days and two nights), two weeks after the exposure initiation. Pair breeding using one male and female per breeding cage was conducted to minimize stress, overcrowding, and other variables. All sires were physically removed from the breeding cages to ensure proper timing of pregnancy. Pb-exposed animals were switched to control water at E10–12 until euthanasia. Study animals were euthanized via CO_2_ asphyxiation followed by double pneumothorax. All animal procedures were approved by the University of Michigan Institutional Animal Care and Use Committee (IACUC). Cardiac puncture was used to collect whole blood from control and Pb-exposed females immediately after euthanasia. Maternal blood Pb levels were measured on an inductively coupled plasma mass spectrometer (ICP-MS) through the Michigan Department of Health and Human Services (*n*=2–3 per group; limit of detection <1.0 μg/dL). The start of animal exposure, mating, removal of sire, switching to control water bottles, and animal euthanasia occurred during the hours of 09:00 to 14:00.

### 2.2. Tissue collection, nucleic acid extraction, and sex determination

The mouse uterus was dissected and placed in a 100mm x 15mm petri dish (Fisher Scientific, #FB0875712) and submerged in 1X PBS, pH 7.4 (Gibco, #10-010-023) on ice. Visual observations including litter size, embryonic viability (determined based on morphological embryonic growth for the given stage), and the number of viable embryos per litter were recorded ^35^. Each embryonic sac and whole embryo were carefully dissected and underwent 1– 2 washes with cold 1X PBS to remove excess blood. The whole embryo tissues were gently blotted on lint-free wipes before recording their weights on a digital weighing scale (Fisher Science Education, #SLF103). All tissues including the embryonic sac and head were collected, labeled according to the embryonic position in the uterus, snap frozen in liquid nitrogen, and placed at –80°C for long-term storage. DNA was extracted from frozen embryonic sac tissues using the QIAamp DNA Blood Mini Kit (Qiagen, #51106) according to the manufacturer’s protocol. DNA isolated from embryonic sacs for each exposure group (total *n*=147) was used to determine the sex of E13–15 embryos using an established protocol via PCR ^36^. A subset of control and Pb-exposed embryonic head samples representing 1 female and 1 male from each litter (using *n*≥9 per sex per group) were used for RNA extraction using the AllPrep DNA/RNA Mini Kit (Qiagen, #80204) supplemented with the RNase-free DNase Set (Qiagen, #79254), according to the manufacturer’s protocol. The nucleic acid concentration and quality was assessed using a NanoDrop 2000 Spectrophotometer (Thermo Scientific, #ND-2000). A total of *n*=38 embryonic heads consisting of *n*=18 control (9 females and 9 males) and *n*=20 Pb-exposed (10 females and 10 males) samples were selected for RNA-seq experiments.

### 2.3. Whole embryo weight analysis

Data for a total of *n*=147 samples were recorded for exposure and sex-specific assessment. The Pearson’s chi-square test was conducted to determine statistical significance between number of viable embryos by exposure group and sex. Whole embryo weight data for *n*=146 representing a total *n*=71 for the control group (*n*=39 females and *n*=32 males) and a total *n*=75 for the Pb-exposed group (*n*=36 females and *n*=40 males) was recorded at animal euthanasia and used for embryo weight assessment by exposure group and sex. Weight data for one control male embryo was excluded from analysis as the data was not recorded during tissue collection. Linear mixed-effects models were used assess the effect of Pb exposure on embryonic weight by accounting for sex as a fixed effect and litter as a random effect. The analysis was performed in R statistical software (v2025.5.1.513) using the *lmer* function per the lmertest package, where test statistic was used to calculate statistical significance, and which is built on the lme4 package ^37,38^. Models were also stratified by sex, accounting for litter as a random effect. Trends were visualized using *ggplot2* in RStudio. Results of *p*<0.05 were considered statistically significant.

### 2.4. RNA quantification, library preparation, and RNA-seq

RNA quantification and library preparation were conducted using previously established protocols, and RNA-seq was executed at the University of Michigan (UM) Advanced Genomics Core ^24,39^. Briefly, the Quant-IT RiboGreen RNA Assay kit (ThermoFisher, #R11490) was used to quantify the RNA concentrations. A plexWell (SeqWell) plate-based approach was used for cDNA preparation, quality control, and library preparation. cDNA was prepared using the seq well cDNA module (SeqWell, #301035), purified using MAGwise Paramagnetic beads (SeqWell, #MG10000), and cDNA concentrations were quantified using Quant-iT PicoGreen dsDNA Assay Kit (ThermoFisher Scientific, #P11496). The resulting fluorescence was measured by the SpectraMax M5e using a pre-set protocol for PicoGreen. The plexWell LP384 Library Preparation Kit (SeqWell, #LP684X) was used to barcode cDNA libraries using approximately 10ng of cDNA input per sample. Barcoded samples were pooled, bead purified and quantified on the Agilent Bioanalyzer (High Sensitivity DNA 5000 Kit). Randomly pooled libraries were sequenced on the Illumina NovaSeq 6000 using 151bp paired-end reads at a sequencing depth of >30 million reads per sample.

### 2.5. Differentially expressed genes based on exposure group

To process the RNA-sequencing data, reads (FASTQ files) were demultiplexed back into individual samples based on individual barcodes. The sequence quality was subsequently determined using FastQC (v 0.12.1) ^40^. Reads were aligned and counted using Salmon (v1.9.0), relative to the Gencode M35 mouse reference genome ^41^. Read count matrices were loaded into R via edgeR (v4.6.3) ^42^. All samples had library sizes >10 million mapped reads for downstream analyses. Lowly expressed genes were filtered by using *filterByExpr*, normalized by using *calcNormFactors*, and the sample dispersion was estimated by using *estimateDisp* (default settings were utilized for each function). Differentially expressed genes by exposure group (comparison between Pb-exposed embryonic head relative to control) were identified in all mice and then stratified by sex. Each gene comparison was calculated using the quasi-likelihood F-test (QLFT) negative binomial generalized linear model (GLM). Genes significant at a false discovery rate (FDR)-adjusted *p*-values (FDR<0.05) were considered significant differentially expressed genes. The data distribution by exposure group (*n*=18 for control and *n*=20 for Pb) and subsequent sex stratification (*n*=9 per sex for control and *n*=10 per sex for Pb) were visualized by multidimensional scaling plots using the plotMDS function ^43^. Differentially expressed genes by exposure group for all tested embryonic heads and for sex-specific models were visualized using volcano plots (supplemented by ggrepel). Differentially expressed genes from the combined sex, and sex-specific model comparisons were used to generate a Venn diagram of differentially expressed gene overlap between sexes. The LogFC by exposure group between females and males were calculated using the Pearson’s correlation test for all genes. Data visualization was conducted using *ggplot2*.

### 2.6. Gene set enrichment analysis based on exposure group

Gene Set Enrichment Analysis was performed using LRpath on Gene Ontology (GO) terms and Kyoto Encyclopedia of Genes and Genomes (KEGG) pathways in both sex-combined, and sex-stratified models ^44^. Enriched GO categories including biological process (GOBP), cellular components (GOCC), and molecular functions (GOMF), as well as KEGG pathways, were identified. A stringent threshold for LRpath analysis (FDR<0.001) was used to select the most relevant pathways associated with Pb exposure. Terms containing more than 1000 genes were excluded. The ten most statistically significant GO terms and KEGG pathways from each of the combined sex and sex-stratified models were selected for visualization.

### 2.7. Differentially expressed imprinted genes, X linked genes, and associated pathways based on exposure group, stratified by sex

The differentially expressed genes by exposure group using the combined sex model and sex-stratified models were further filtered by a list of known 300 gametic and non-canonical imprinted genes including those from the geneimprint database ^25^. An X-linked gene list was generated using the UCSC Genome Browser mouse assembly (GRCm39/mm39; Gencode VM35) for the X chromosome (chrX: 1–169,476,592). All differentially expressed imprinted genes and differentially expressed X-linked genes upon Pb exposure were confirmed by using the log_2_ counts per million changes calculated using the *cpm* function of normalized data. The resulting imprinted genes and X-linked genes from the combined sex, F, and male data comparisons were used to generate a Venn diagram of overlapping genes between sexes. A comparison of the LogFC by exposure group between females and males were calculated using the Pearson’s correlation test. A heatmap of differentially expressed imprinted genes within significantly enriched GOBP terms or KEGG pathways was generated using ComplexHeatmap (v2.24.1) ^45^. Gene expression was normalized with the *cpm* function in edgeR (v4.6.3). For genes present in multiple enriched categories, annotation was assigned to the category containing the largest number of differentially expressed imprinted or X-linked genes.

### 2.8. Differentially expressed genes and associated pathways based on sex, stratified by exposure group to identify sexually dimorphic autosomal genes

Sexually dimorphic differentially expressed genes were assessed by sex using a comparison between male embryonic heads relative to F, exclusively for the control group. Data distribution by sex for each exposure group (*n*=9 females and 9 males for control; n=10 females and 10 males for Pb) was visualized using the MDS plots, after filtering and normalizing for low read counts using edgeR ^42^. Each sex-based comparison was calculated using the QLFT negative binomial GLM. The sexually dimorphic genes were compared against the differentially expressed genes identified based on the exposure for the combined model, stratified by sex to generate the Venn diagrams. Over-representation analyses of sexually dimorphic Pb-sensitive differentially expressed genes were conducted using the *enrichGO* and *enrichKEGG* functions in clusterProfiler (v4.16.0) ^46^. Enrichment was determined at FDR<0.05. Fisher’s exact tests were applied to evaluate overlaps between sexually dimorphic and Pb-sensitive differentially expressed genes using the *fisher.test* function in stats (v4.5.1).

## 3. RESULTS

### 3.1. Pb exposure during preconception and gestation increases embryonic weight

Adult mice were exposed to either control or Pb in drinking water at 32 ppm two weeks before mating until E10–12 and euthanized at E13–15 at mid-gestation. The average maternal blood Pb level of 9.7 μg/dL was detected for the Pb-exposed group, while the control group was below the limit of detection. This is a human-relevant exposure level and was measured after 4 weeks of exposure to experimental water and food provided *ad libitum* ^47^. The control group produced 7.89 viable embryos per litter from 9 successful pregnancies at 89.87% viability, while 7.60 viable embryos per litter were produced from 10 successful pregnancies at 90.48% viability for the Pb group (**Table 1** and **Supplementary** Figure 1). No significant differences were observed in litter sizes and the number of viable embryos between exposure group (**Supplementary** Figure 1). The sex distribution did not significantly differ between the control (39 females and 32 males) and the Pb-exposed groups (36 females and 40 males) (χ² (1, N=147) = 0.56, *p*=0.45).

**Table 1.** Summary of Litter Sizes and Embryo Viability.

| Group | Number of litters | Total number of embryos | Total number of viable embryos | Percent viability |
| --- | --- | --- | --- | --- |
| Control | 9 | 79 | 39 (F); 32 (M) = 71 | 89.87% |
| Pb | 10 | 84 | 36 (F); 40 (M) = 76 | 90.48% |

Whole embryo weights at E13–15 were recorded to evaluate embryonic weight differences by exposure group and sex. Weight records for each embryo by litter, exposure group, and sex were analyzed for *n*=146. The raw data show a trend toward higher absolute embryonic weights 0.11–0.42g for the Pb-exposed group relative to the control group 0.08–0.29g (**Supplementary Figure S2**). This trend was observed in both sexes (**Supplementary Figure S3**) for each exposure group (control F: 0.08–0.26g and M: 0.08–0.29g; Pb-exposed F: 0.11–0.42g and M: 0.12–0.41g).The linear mixed-effects model for exposure group (**Figure 2A**), controlling for litter effects showed that Pb exposure had a significant effect on embryonic weight, relative to control (*p*=0.0068). Pb-exposed whole embryos (average weight at 0.24g) had significantly higher total weights compared to control (average weight at 0.16g), independent of sex and litter. A separate linear mixed-effects model was used to analyze the Pb exposure effects on embryonic weights stratified by sex, while controlling for litter effects (**Figure 2B**). The Pb group displayed a significantly higher (*p*=0.00793) embryonic weights for females (average weight at 0.25g), relative to the control females (average weight at 0.15g). Similarly, the Pb-exposed male embryonic weights (average weight at 0.24g) were significantly higher (*p*=0.00746), relative to control males (average weight at 0.17g). Collectively, these results suggest that the Pb-exposed whole embryos weigh significantly more than controls, independent of sex and litter, upon preconception and gestational exposure to Pb.

**Figure 1.**
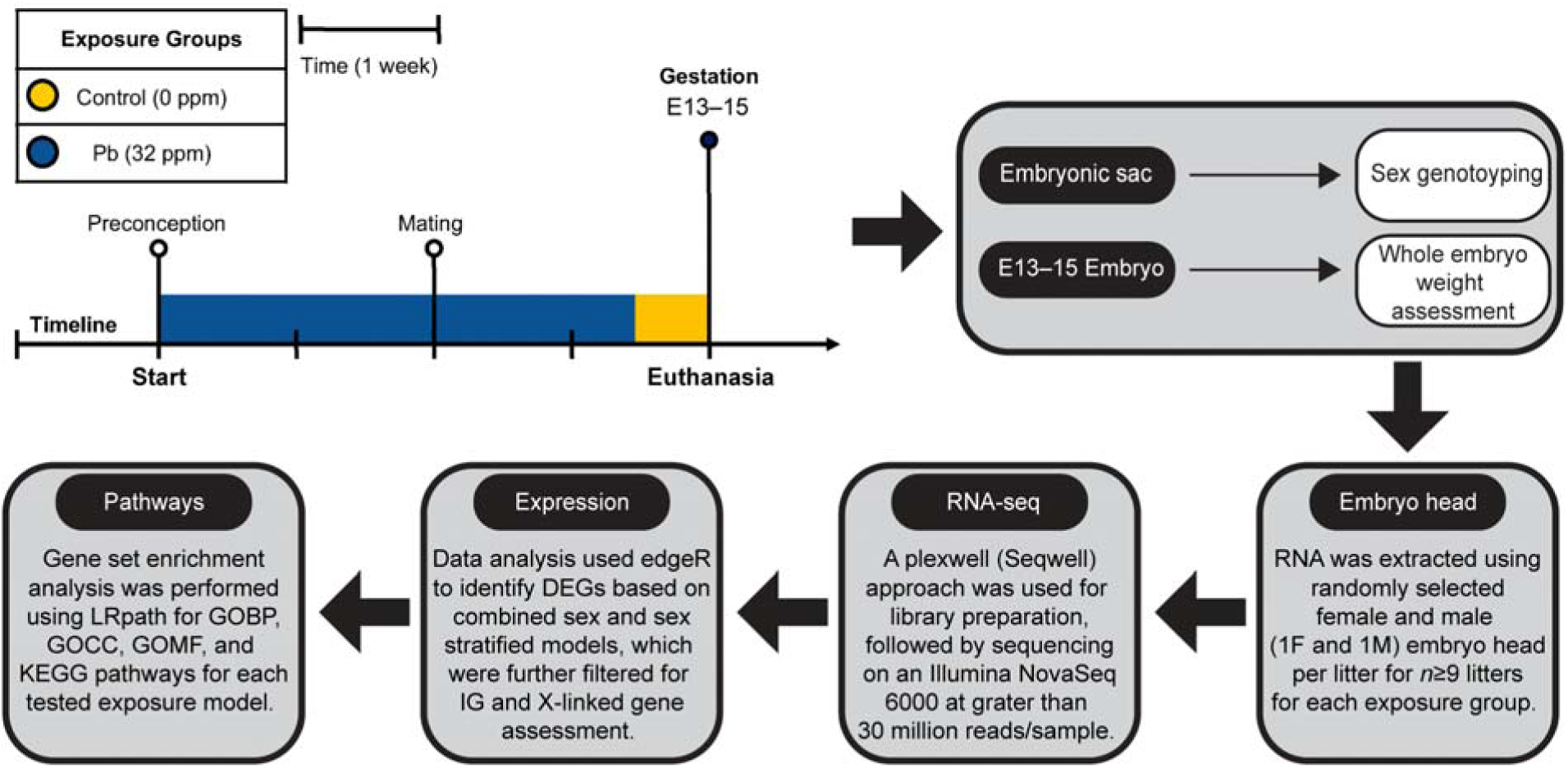
Experimental design for the perinatal Pb exposure study. C57BL/6J adult females at 10–12 weeks of age were subjected to either control (yellow) or Pb-spiked water at 32 ppm (dark blue) starting 2 weeks before mating, continued until embryonic day (E)13–15. All animals were time-mated with their male counterparts for two nights at two weeks after the start of exposure. Pb-exposed animals were switched to control water at E10–12 prior to euthanasia. The whole embryo tissues were collected at E13–15 and weighted prior to embryonic head collection. DNA was extracted from the embryonic sac for sex genotyping. A female (F) and male (M) embryo head RNA were extracted from each litter (n=9 for control and n=10 for the Pb-exposed group), to account for litter effects. RNA was converted to cDNA and libraries were prepared for sequencing, which was subsequently used for gene expression data analysis using edgeR and gene set enrichment assessment using LRpath-based methods. DEGs: differentially expressed genes; IG: imprinted gene.

**Figure 2.**
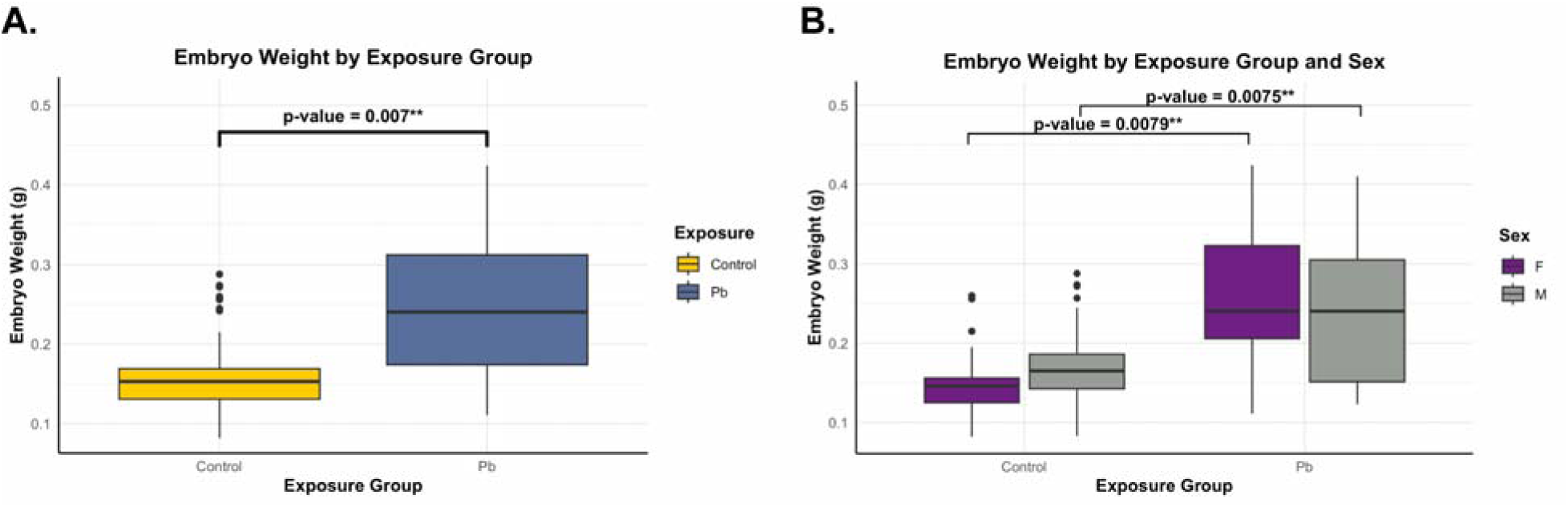
Whole embryo weight assessment by exposure group and sex. The data represent *n*=146 total embryo weights representing all animals used for the study; *n*=1 whole embryo weight (control male) was excluded from analysis. Statistical significance is shown as *p*<0.01 (**). (**A**) Linear mixed effects model for the whole embryo weight by exposure group. The box plot represents the combined sex model indicating control (yellow) and Pb (dark blue) exposure groups that were assessed using the exposure group as a fixed effect and litter as a random effect. (**B**) Linear mixed effects model for the whole embryo weight by exposure group and sex. The box plot represents the model accounting for exposure group and sex, indicating females (F; purple) and males (M; grey) for control and Pb exposure groups that were assessed using exposure group and sex as a fixed effect, and litter as a random effect.

### 3.2. Perinatal Pb exposure leads to differential neuronal gene expression in the embryonic head

Bulk RNA-seq was conducted to determine the overall gene expression changes between control and Pb-exposed groups, using 1 female and 1 male representative embryonic heads per litter and selected at random (control: *n*=9 females and 9 males; Pb: *n*=10 females and 10 males). The final library sizes after filtering for lowly expressed genes and normalizing ranged between 14,032,468–39,354,980, with 22,432,193 average reads per sample (**Supplementary Table S1**), indicating consistent sequencing depth between biological replicates within each exposure and sex group. The MDS plots indicate that sample clustering for the overall transcriptomic data is driven mainly by exposure group rather than by sex (**Supplementary Figure S4**). A total of 26,937 genes were detected, with 15,372 genes remaining after filtering and normalization (**Table 2**). The magnitude of gene expression changes by exposure for the combined sex and sex-stratified models are shown with volcano plots in **Figures 3–5** and **Supplementary Figure S5.**

**Figure 3.**
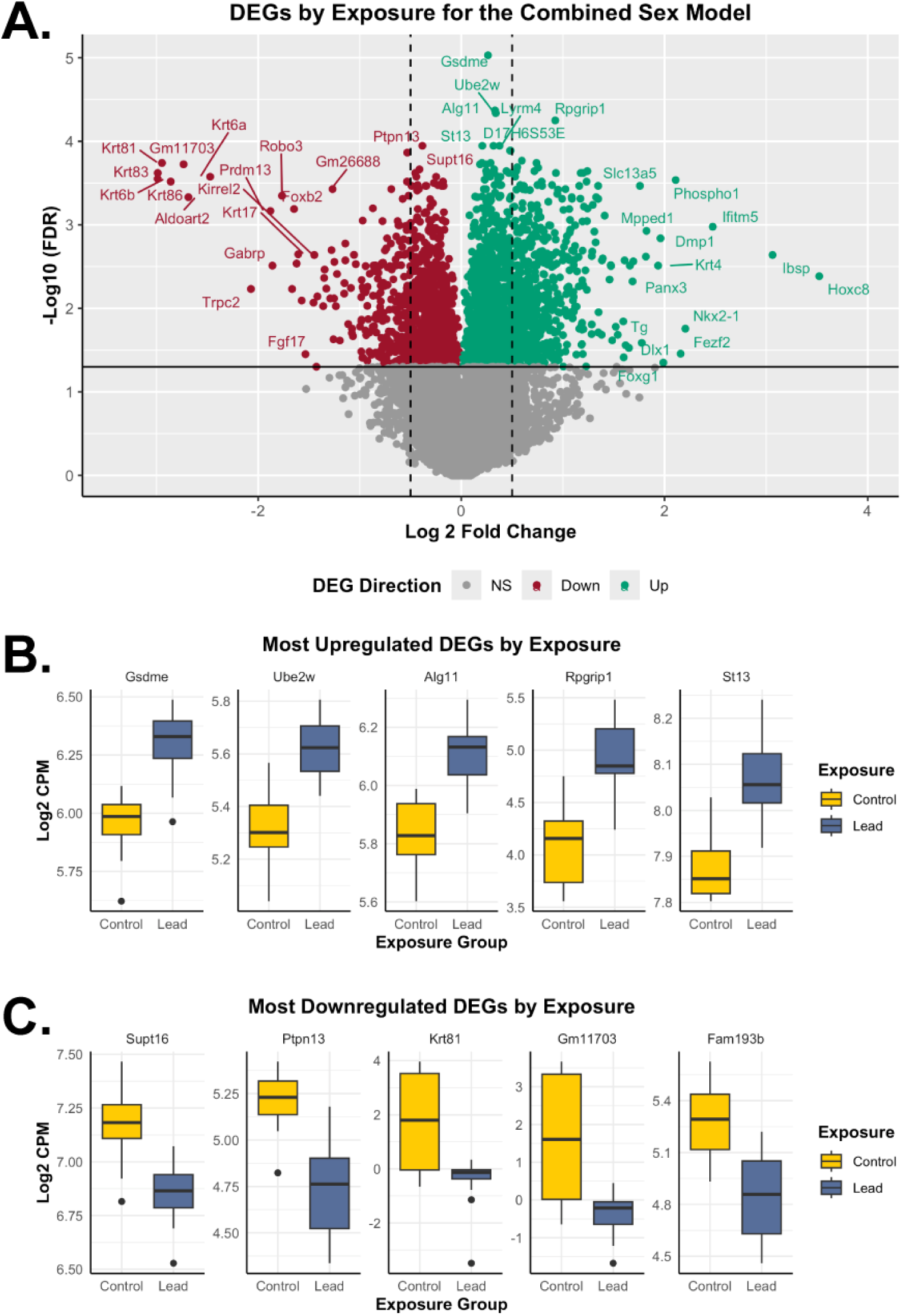
Differentially expressed genes based on exposure group for the combined sex model. The combined model shows results related to *n*=38 RNA-seq samples, including *n*=18 for control and *n*=20 for the Pb exposure groups. (**A**) The volcano plot represents upregulated (teal; LogFC>0; FDR<0.05) and downregulated (red; LogFC<0; FDR<0.05) differentially expressed genes by Pb exposure relative to control. The dotted lines indicate 0.5 LogFC, while non-significant (NS) genes are shown in grey. (**B**) The top 5 significantly upregulated differentially expressed genes (based on FDR) by exposure group. Gene expression of *Gsdme*, *Ube2w*, *Alg11*, *Rpgrip1*, and *St13* is represented by boxplots, based on control (yellow) and Pb (dark blue) exposure groups. (**C**) The top 5 significantly downregulated differentially expressed genes (based on FDR) by exposure group. Gene expression of *Supt16*, *Ptpn13*, *Krt81*, *Gm11703*, and *Fam193b* is represented by boxplots, based on control (yellow) and Pb (dark blue) exposure groups. DEGs: differentially expressed genes.

**Figure 4.**
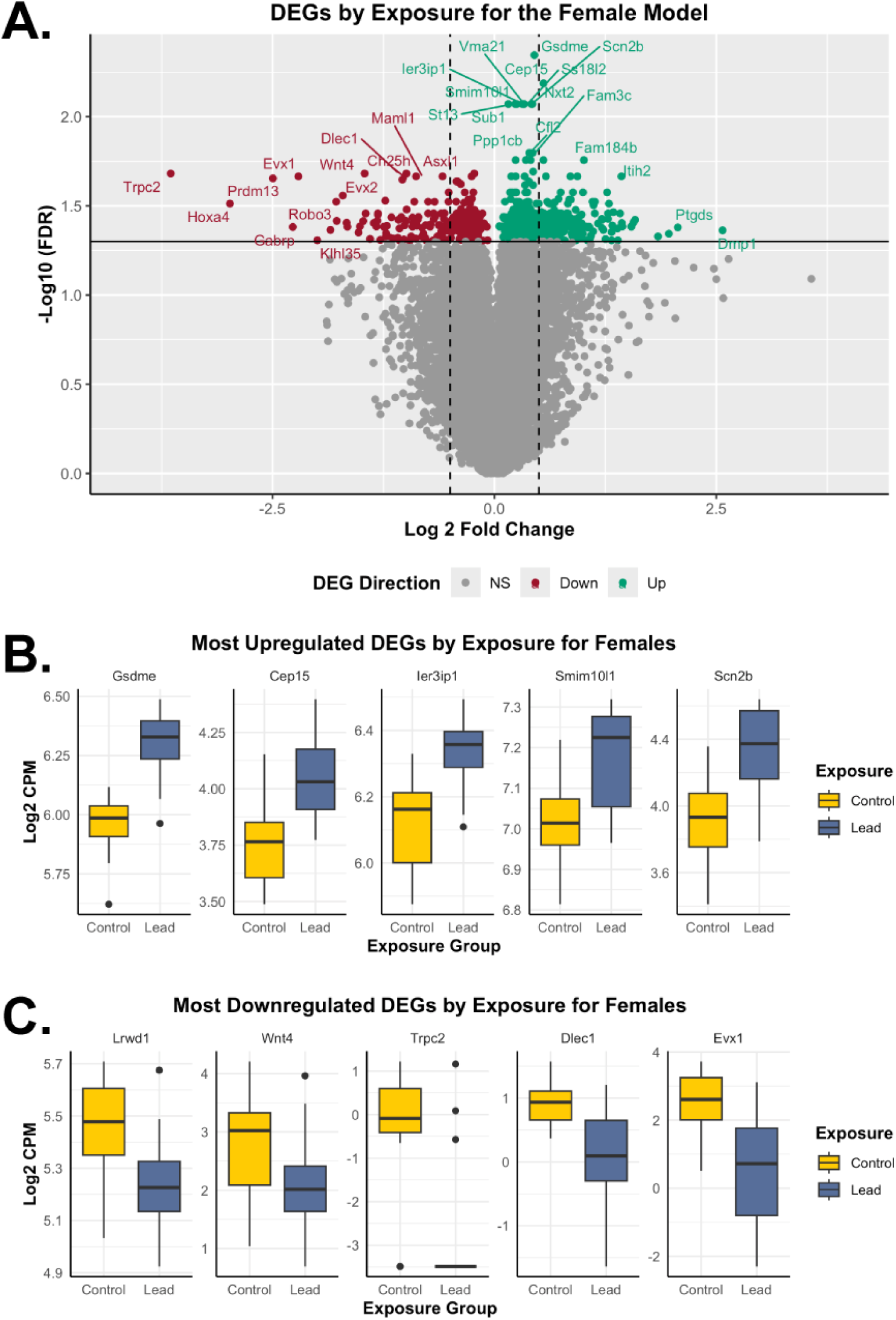
Differentially expressed genes based on exposure group for the female model. The female model shows results related to *n*=19 RNA-seq samples, including *n*=9 for control and *n*=10 for the Pb exposure groups. (**A**) The volcano plot represents upregulated (teal; LogFC>0; FDR<0.05) and downregulated (red; LogFC<0; FDR<0.05) differentially expressed genes in Pb-exposed females, relative to control females. The dotted lines indicate 0.5 LogFC, while non-significant (NS) genes are shown in grey. (**B**) The top 5 significantly upregulated differentially expressed genes (based on FDR) by exposure group for females. Gene expression of *Gsdme*, *Cep15*, *Ier3ip1*, *Smim10l1*, and *Scn2b* is represented by boxplots, based on control (yellow) and Pb (dark blue) exposure groups. (**C**) The top 5 significantly downregulated differentially expressed genes (based on FDR) by exposure group for females. Gene expression of *Lrwd1*, *Wnt4*, *Trpc2*, *Dlec1*, and *Evx1* is represented by boxplots, based on control (yellow) and Pb (dark blue) exposure groups. DEGs: differentially expressed genes.

**Figure 5.**
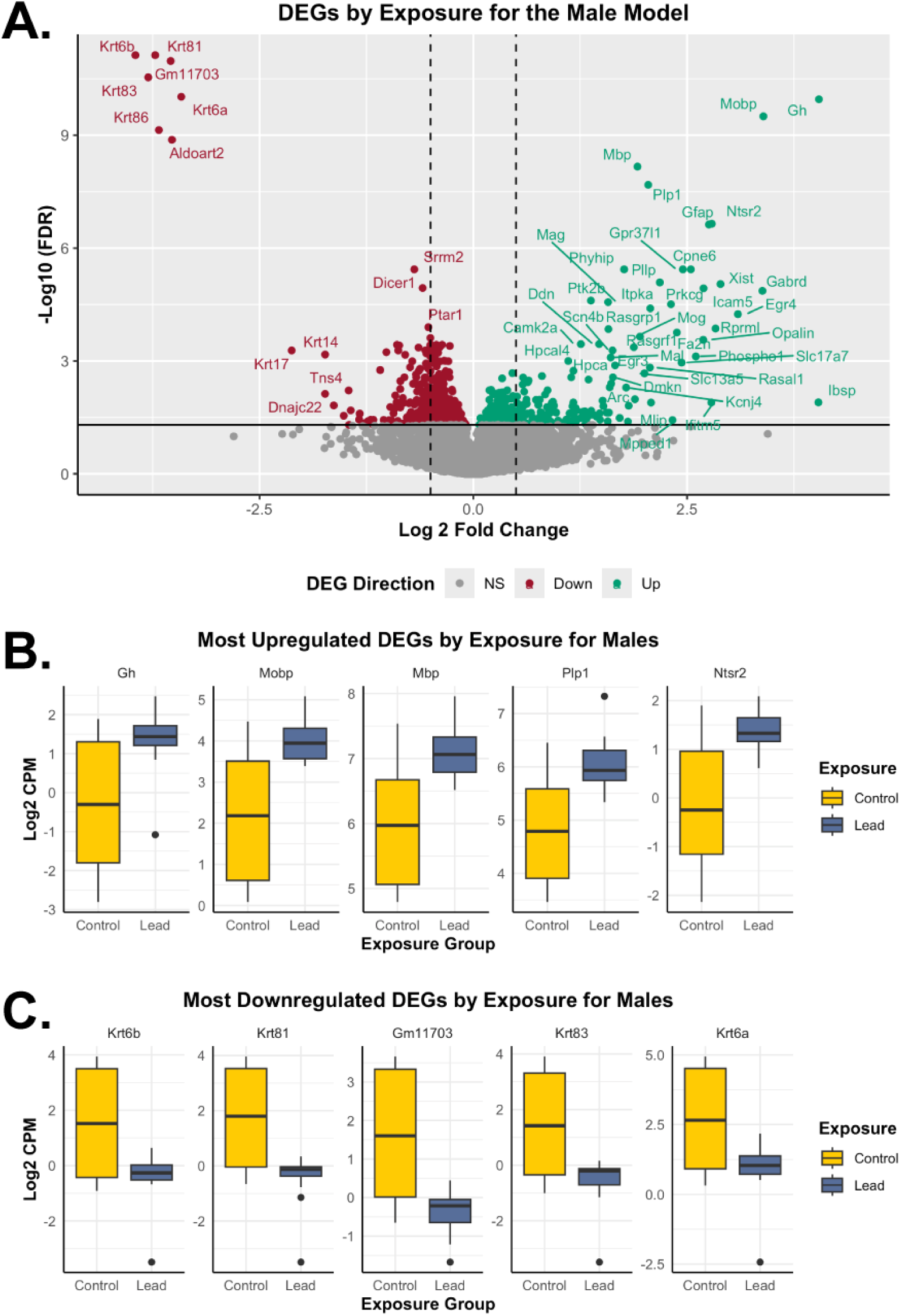
Differentially expressed genes based on exposure group for the male model. The male model shows results related to *n*=19 RNA-seq samples, including *n*=9 for control and *n*=10 for the Pb exposure groups. (**A**) The volcano plot represents upregulated (teal; LogFC>0; FDR<0.05) and downregulated (red; LogFC<0; FDR<0.05) differentially expressed genes in Pb-exposed males, relative to control males. The dotted lines indicate 0.5 LogFC, while non-significant (NS) genes are shown in grey. (**B**) The top 5 significantly upregulated differentially expressed genes (based on FDR) by exposure group for males. Gene expression of *Gh*, *Mobp*, *Mbp*, *Plp1*, and *Ntsr2* is represented by boxplots, based on control (yellow) and Pb (dark blue) exposure groups. (**C**) The top 5 significantly downregulated differentially expressed genes (based on FDR) by exposure group for males. Gene expression of *Krt6b*, *Krt81*, *Gm11703*, *Krt83*, and *Krt6a* is represented by boxplots, based on control (yellow) and Pb (dark blue) exposure groups. DEGs: differentially expressed genes.

**Table 2.** Number of Differentially Expressed Genes Based on Exposure and Sex for the Embryonic Head at E13–15.

| Category | Total number of genes sequenced | Total number of genes after filtering and normalization | Number of DEGs with $\vert \text{LogFC}\vert > 0$; $\text{FDR} < 0.05^*$ | Number of DEGs upregulated with Pb ($\text{LogFC} > 0$; $\text{FDR} < 0.05$) | Number of DEGs downregulated with Pb ($\text{LogFC} < 0$; $\text{FDR} < 0.05$) |
| --- | --- | --- | --- | --- | --- |
| Combined sex | 26,937 | 15,372 | 2,662 | 1,551 | 1,111 |
| Females |  |  | 668 | 446 | 222 |
| Males |  |  | 709 | 306 | 403 |

\* $\text{LogFC}$ : log fold change; $\text{FDR}$ : false discovery rate.

For the combined sex model, 17.3% (2,662) of total sequenced genes were differentially expressed genes upon exposure to Pb, relative to the control group (**Table 2** and **Supplementary Table S2**). Approximately 58.3% (1,551) and 41.7% (1,111) of the combined sex differentially expressed genes were significantly increased and decreased upon Pb exposure, respectively (**Figure 3A**). For Pb exposure in the combined model, the most significantly upregulated differentially expressed genes included *Gsdme*, *Ube2w*, *Alg11*, *Rpgrip1*, and *St13* (**Figure 3B**), while the most downregulated differentially expressed genes contained *Supt16*, *Ptpn13*, *Krt81*, *Gm11703*, and *Fam193b* (**Figure 3C**).

Sexually dimorphic effects were shown in males for *Krt81* and *Gm11703* (**Supplementary** Figure 5A). The sex-stratified assessment for the female model indicated that 4.4% (668) of total sequenced genes were dysregulated upon Pb exposure (**Table 2** and **Supplementary Table S3**). Approximately 66.8% (446) and 33.2% (222) of the differentially expressed genes for the female model were significantly increased and decreased upon Pb exposure, respectively (**Figure 4A**). The most significantly upregulated differentially expressed genes for the female model included *Gsdme*, *Cep15*, *Ier3ip1*, *Smim10l1*, and *Scn2b* (**Figure 4B**), while the most downregulated differentially expressed genes included *Lrwd1*, *Wnt4*, *Trpc2*, *Dlec1*, and *Evx1* (**Figure 4C**) by Pb exposure, with sex-specific gene expression comparisons to the combined sex model (**Supplementary** Figure 5B). For males, 4.61% (709) of total captured genes were dysregulated upon Pb exposure (**Table 2** and **Supplementary Table S4**). Approximately 43.2% (306) and 56.8% (403) of the differentially expressed genes among males were significantly increased and decreased upon Pb exposure, respectively (**Figure 5A**). The most significantly upregulated differentially expressed genes among males included *Gh*, *Mobp*, *Mbp*, *Plp1*, and *Ntsr2* (**Figure 5B**), while the most downregulated differentially expressed genes included *Krt6b*, *Krt81*, *Gm11703*, *Krt83*, and *Krt6a* (**Figure 5C**), showing sexually dimorphic effects of Pb exposure (**Supplementary** Figure 5C).

Taken together, 3.5% (103) of total differentially expressed genes were found common between all three tested models, with 3.8% (112) and 4.9% (144) found exclusively in the female and male models, respectively (**Supplementary Figure S6**). A statistically significant positive correlation (r=0.52, *p*<0.001) was observed between the LogFCs between the two sexes (**Supplementary Figure S7**). These results indicate that Pb exposure significantly dysregulates gene expression with some effects in common and some sex-specific effects.

### 3.3. Pb exposure dysregulates neurodevelopmental pathways

To assess the biological pathways and disease mechanisms impacted by gene expression changes upon exposure and sex during preconception and gestational Pb exposure, Gene Ontology and KEGG enrichment analyses were conducted using LRpath (**Figures 6** and **7**, **Supplementary Tables S5–S10**). Comparison of the combined sex and sex-stratified models showed that GOBP (total 249), GOCC (total 91), GOMF (total 87), and KEGG (total 74) pathways were significantly impacted by gestational exposure to Pb (**Figure 6**). For GOBP, 23.7% (59) of common biological pathways were enriched between all three tested models, while 16.5% (41) and 18.9% (47) were enriched exclusively in the female and male models, respectively. The most significantly dysregulated GOBP pathways upon Pb exposure included synaptic signaling, embryonic morphogenesis, epigenetic regulation of gene expression, among others (**Supplementary Tables S5**, **S7**, and **S9**). Pb exposure significantly downregulated the GOBP DNA metabolic process in the female model, while DNA damage response, sexual reproduction, and keratinization processes were significantly downregulated in the male model (**Figure 7**).

**Figure 6.**
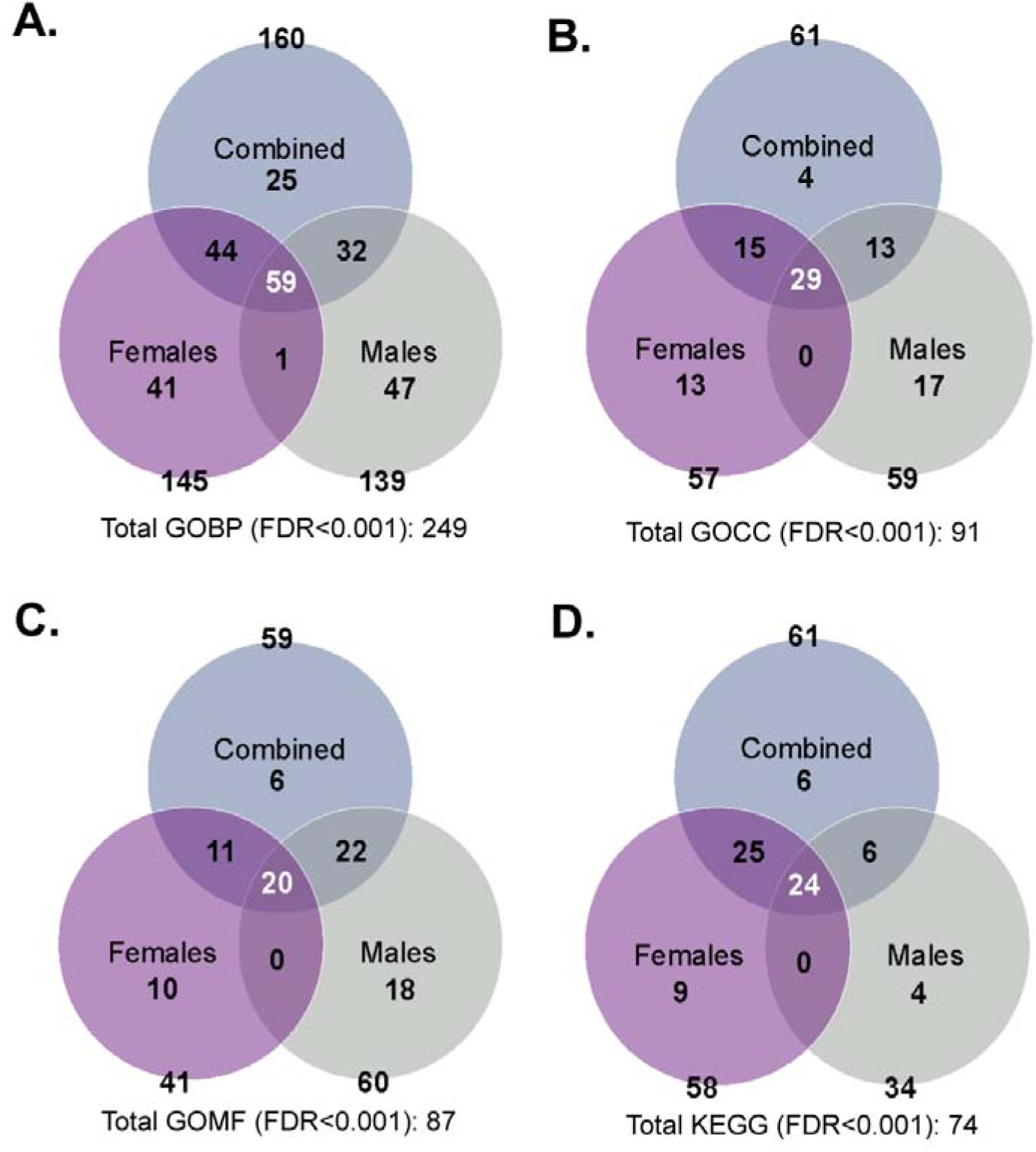
Overview of the most significant gene ontology and KEGG pathways disrupted by gestational Pb exposure. The Venn diagrams represent data related to the combined sex model (blue), female model (purple), and male model (grey) for the most significant pathways impacted by Pb exposure (FDR<0.001). (**A**) Gene Ontology Biological processes (GOBP) impacted by exposure and sex. (**B**) Gene Ontology Cellular components (GOCC) impacted by exposure and sex. (**C**) Gene Ontology Molecular functions (GOMF) impacted by exposure and sex. (**D**) KEGG pathways impacted by exposure and sex. Collectively, 249 GOBP, 91 GOCC, 87 GOMF, and 74 KEGG pathways are dysregulated by gestational Pb exposure in the embryonic head, based on the three models assessed.

**Figure 7.**
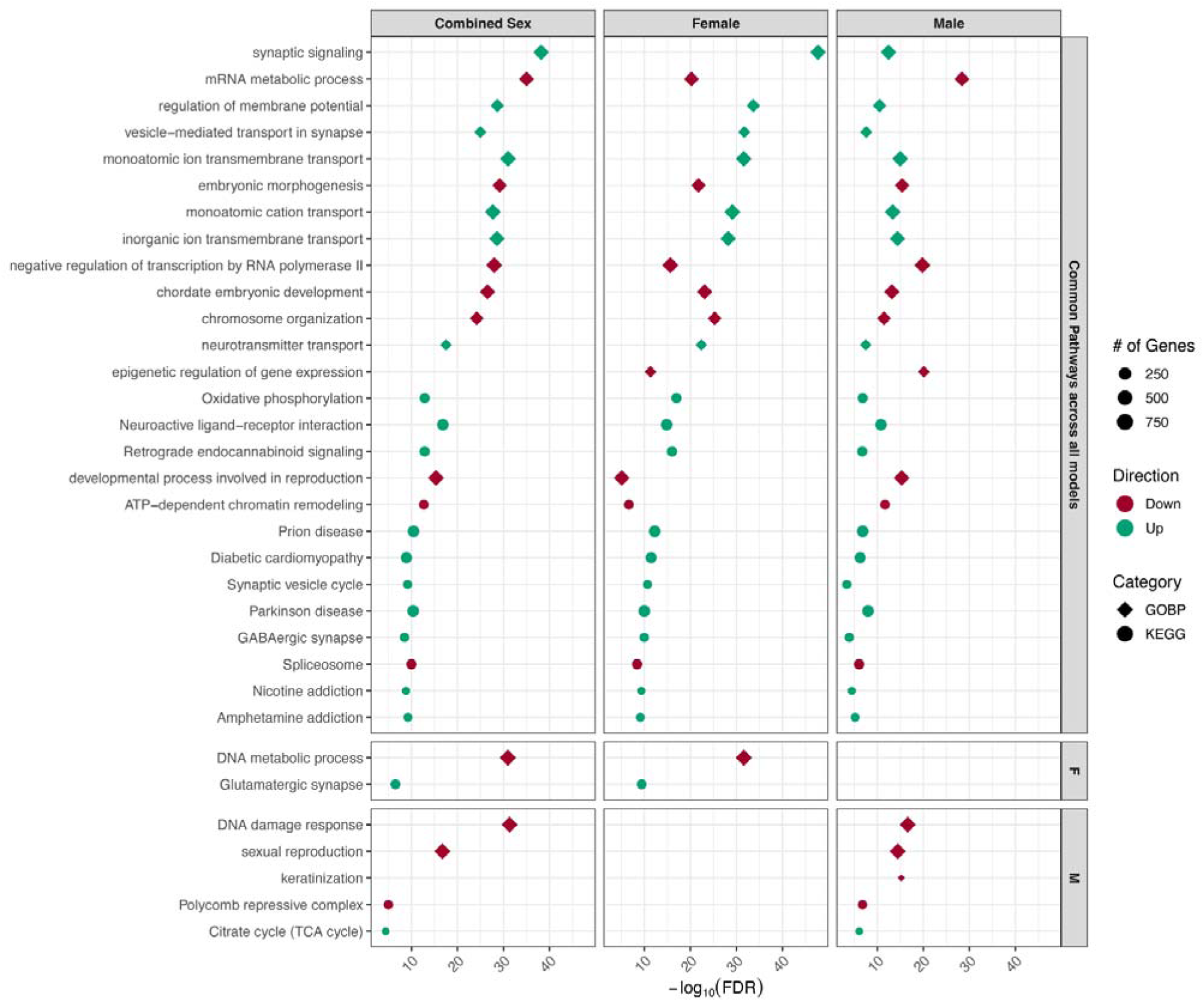
The most enriched biological processes and KEGG pathway terms associated with gestational Pb exposure. Results are shown for combined sex, female, and male models (top subheadings) with gene ontology biological process (GOBP; diamond) and Kyoto Encyclopedia of Genes and Genomes (KEGG; circle). The model-specific GOBP and KEGG comparisons are listed in subheadings on the right. Pathway enrichment direction is indicated by teal (upregulated) and red (downregulated). Point size reflects the number of genes contributing to enrichment.

For GOCC, 31.9% (29) of common cellular components were enriched between all three tested models, while 14.3% (13) and 18.7% (17) were enriched exclusively in the female and male models, respectively. The most significantly dysregulated GOCC upon Pb exposure included chromatin, presynapse, and transmembrane transporter complex, among others (**Supplementary Tables S5**, **S7**, and **S9**). For GOMF, 23.0% (20) of common molecular functions were enriched between all three tested models, while 11.5% (10) and 20.7% (18) were enriched exclusively in the female and male models, respectively. The most significantly dysregulated GOMF upon Pb exposure were chromatin binding, cis-regulatory region seq-specific DNA binding, neurotransmitter receptor activity, and modification-dependent protein binding, among others (**Supplementary Tables S5**, **S7**, and **S9**).

For KEGG, 32.4% (24) of common disease pathways were enriched between all three tested models, while 12.2% (9) and 24.3% (4) were enriched exclusively in the female and male models, respectively. The most significantly dysregulated KEGG pathways upon Pb exposure included oxidative phosphorylation, neuroactive ligand-receptor interaction, Parkinson’s disease, and diabetic cardiomyopathy, among others (**Supplementary Tables S6**, **S8**, and **S10**). The KEGG glutamatergic synapse was significantly upregulated upon Pb exposure in the female model (**Figure 7**). Similarly, the Polycomb repressive complex (PRC) and the tricarboxylic acid (TCA) cycle were significantly down and upregulated, respectively, upon Pb exposure in the male model (**Figure 7**). Collectively, these findings reveal sex-specific dysregulation of neuronal and metabolic pathways in the developing brain following preconception and gestational exposure to Pb that contribute to disease risk.

### 3.4. Pb exposure dysregulates imprinted genes in a sex-specific manner

Given the role of imprinted genes in fetal growth and regulation, a list of 300 canonical and non-canonical imprinted genes were used to filter differentially expressed genes by exposure for the combined sex and sex-stratified models (**Supplementary Tables S11–S13**). A total of 161 imprinted genes were captured after filtering (**Table 3**). The magnitude of imprinted gene expression changes by exposure for the combined sex and sex-stratified models are shown with volcano plots in **Supplementary Figures S8–S10**.

**Table 3.**
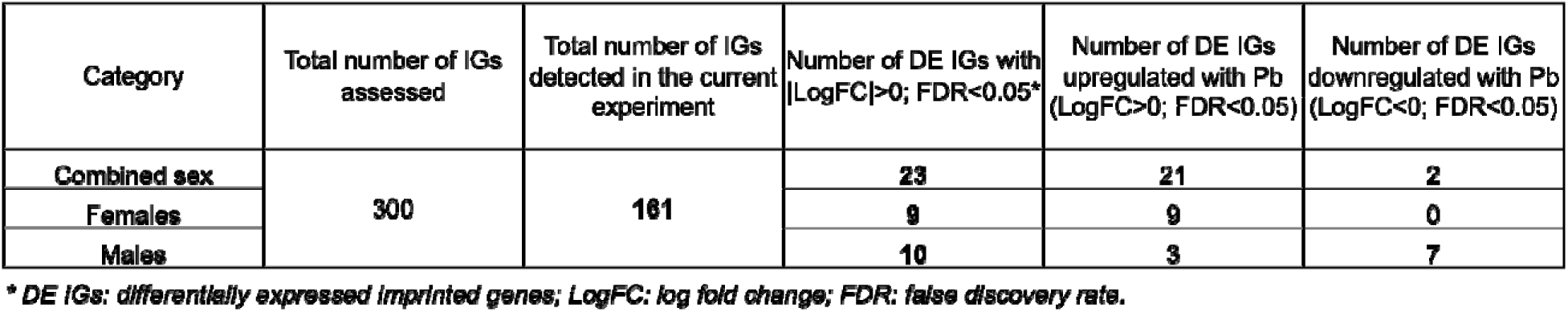
Number of Differentially Expressed Imprinted Genes Based on Exposure and Sex for the Embryonic Head at E13–15.

| Category | Total number of IGs assessed | Total number of IGs detected in the current experiment | Number of DE IGs with $\vert \text{LogFC}\vert > 0$; $\text{FDR} < 0.05^*$ | Number of DE IGs upregulated with Pb ($\text{LogFC} > 0$; $\text{FDR} < 0.05$) | Number of DE IGs downregulated with Pb ($\text{LogFC} < 0$; $\text{FDR} < 0.05$) |
| --- | --- | --- | --- | --- | --- |
| Combined sex | 300 | 161 | 23 | 21 | 2 |
| Females |  |  | 9 | 9 | 0 |
| Males |  |  | 10 | 3 | 7 |

\* DE IGs: differentially expressed imprinted genes; $\text{LogFC}$ : log fold change; $\text{FDR}$ : false discovery rate.

For the combined sex model, 23 (14.3%) of total captured imprinted genes were differentially expressed upon Pb exposure relative to the control group (**Table 3** and **Supplementary Table S11**). Approximately 21 (91.3%) and 2 (8.7%) of the combined sex differentially expressed imprinted genes were significantly increased and decreased upon Pb exposure, respectively (**Figure 8**). The most upregulated differentially expressed imprinted genes for combined sex included *Ifitm10*, *Rasgrf1*, *Qpct*, *Slc22a2*, and *Snrpn*, while the most downregulated differentially expressed imprinted genes included *Etv6*, and *Id1* (**Figure 8B** and **Supplementary Figure S8**).

**Figure 8.**
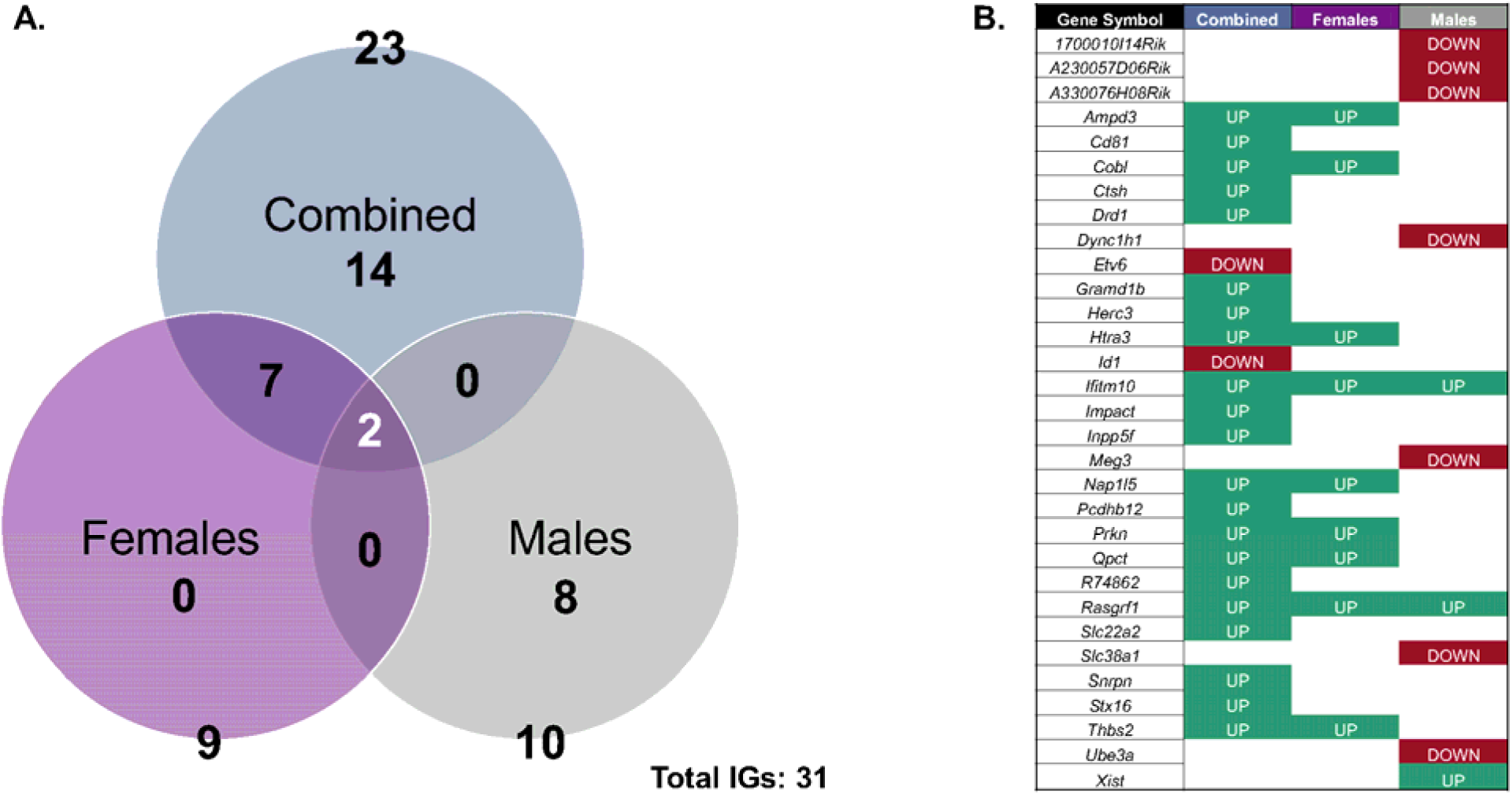
Combined sex, female, and male model-specific differentially expressed imprinted gene comparisons. (**A**) Venn diagram of differentially expressed imprinted gene comparisons between each tested model. The combined model shows 23 differentially expressed imprinted genes related to *n*=38 RNA-seq samples, including *n*=18 for control and *n*=20 for the Pb exposure groups. The female/male models show 9/10 differentially expressed imprinted genes, respectively, related to *n*=19 RNA-seq samples, including *n*=9 for control and *n*=10 for the Pb exposure groups. Collectively, 31 differentially expressed imprinted genes were dysregulated by gestational exposure to Pb in the embryonic head. (**B**) Complete list of differentially expressed imprinted genes is recorded based on directionality of gene expression changes. Upregulated (teal; LogFC>0; FDR<0.05) and downregulated (red; LogFC<0; FDR<0.05) differentially expressed imprinted genes upon Pb exposure relative to control are indicated in respective boxes, comparing the three tested models. IGs: imprinted genes.

Among females, 9 (5.6%) of total captured imprinted genes were dysregulated upon exposure to Pb, with 100% of the differentially expressed imprinted genes significantly upregulated upon Pb exposure (**Table 3** and **Supplementary Table S12**). The most significantly differentially expressed imprinted genes by Pb exposure for the female model included *Ifitm10*, *Htra3, Rasgrf1*, *Nap1l5*, and *Qpct* (**Figure 8B** and **Supplementary Figure S9**). Among males, 10 (6.2%) of total captured imprinted genes were dysregulated upon Pb exposure (**Table 3** and **Supplementary Table S13**). Three of these imprinted genes were upregulated and seven were downregulated with Pb exposure (**Figure 8B**). The most significantly upregulated differentially expressed imprinted genes for the male model included *Xist* and *Rasgrf1,* while the most downregulated differentially expressed imprinted genes contained *Ube3a*, *Meg3*, and *Slc38a1* (**Supplementary Table S13**), with *Ube3a* showing pronounced sexually dimorphic effects from Pb exposure (**Supplementary Figure S10**).

In summary, 2 (6.5%) of the total differentially expressed imprinted genes were common between all three tested models, and 8 (25.8%) were found exclusively among males (**Figure 8A**). A statistically significant moderate positive correlation (r=0.40, *p*<0.001) was observed between the LogFCs of female vs. male models, indicating a tendency for concordance in differentially expressed imprinted gene expression between sexes (**Supplementary Figure S11**). These results indicate that Pb exposure significantly dysregulates imprinted genes, with some observed sex-specific effects. The overall differentially expressed imprinted genes by exposure group and sex are enriched in GOBP and KEGG pathways (**Figure 9**) responsible for metabolism and biosynthetic processes (amide and organophosphate metabolic processes), gene regulation (negative regulation of transcription), cell division, transport and localization (import into cell and localization within membrane), phagosome, sexual reproduction, and nervous system/diseases (synaptic signaling and behavior).

**Figure 9.**
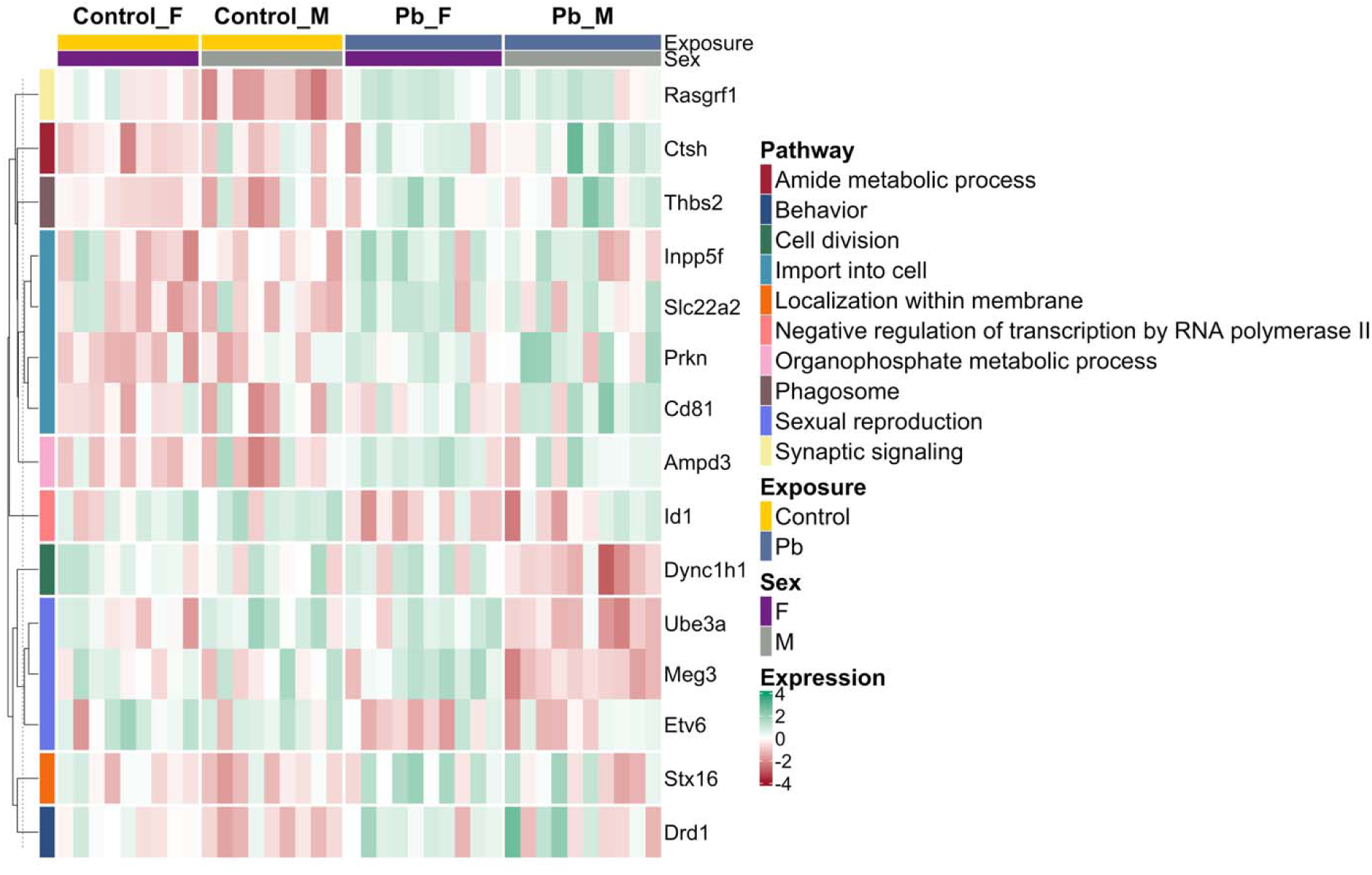
Heatmap of differentially expressed imprinted gene expression within enriched GOBP and KEGG pathways in combined sex and sex-stratified models of Pb exposure. Pathways are indicated by the colored left annotation bar, while exposure (Control: yellow; Pb: dark blue) and sex (Female: F; Male: M) are shown in the top annotation bar. Gene expression levels are represented as z-score scaled log counts per million (logCPM), with gene names displayed on the right.

### 3.5. Pb exposure dysregulates X-linked genes in a sex-specific manner

Given the observed sex-stratified impact on Pb toxicity (**Supplementary Figure S5**), a list of 1,529 X-linked genes was used to filter differentially expressed genes by exposure for the combined sex and sex-stratified models (**Supplementary Tables S14–S16**). A total of 535 X-linked genes were captured after filtering (**Table 4**). The magnitude of X-linked gene expression changes by exposure for the combined sex and sex-stratified models are shown using volcano plots (**Supplementary Figures S12–S14**).

**Table 4.** Number of Differentially Expressed X-linked Genes Based on Exposure and Sex for the Embryonic Head at E13–15.

| Category | Total number of X-linked genes assessed | Total number of X-linked genes detected in the current experiment | Number of DE X-linked genes with $\vert \text{LogFC}\vert > 0$; $\text{FDR} < 0.05^*$ | Number of DE X-linked genes upregulated with Pb ($\text{LogFC} > 0$; $\text{FDR} < 0.05$) | Number of DE X-linked genes downregulated with Pb ($\text{LogFC} < 0$; $\text{FDR} < 0.05$) |
| --- | --- | --- | --- | --- | --- |
| Combined sex | 1,529 | 536 | 107 | 79 | 28 |
| Females |  |  | 27 | 22 | 5 |
| Males |  |  | 24 | 10 | 14 |

\* DE: differentially expressed; $\text{LogFC}$ : log fold change; $\text{FDR}$ : false discovery rate.

For the combined sex model, 107 (20.0%) of total sequenced X-linked genes were differentially expressed upon Pb exposure relative to the control group (**Table 4** and **Supplementary Table S14**). Approximately 79 (73.8%) and 28 (26.1%) of the combined sex differentially expressed X-linked genes were significantly increased and decreased upon Pb exposure, respectively (**Figure 10**). The most significantly upregulated differentially expressed X-linked genes for combined sex included *Tspan7*, *Dynlt3*, *Gabra3*, *Tceal1*, and *Nxt2*, while the most downregulated differentially expressed X-linked genes included *Med12*, *Bcor*, *Brwd3*, *Huwe1*, and *Hcfc1* (**Figure 10B** and **Supplementary Figure S12**). The sex stratified assessment for females indicated that 27 (5.1%) of total sequenced X-linked genes were dysregulated upon Pb exposure (**Table 4** and **Supplementary Table S15**). Approximately 22 (81.9%) and 5 (18.5%) of the differentially expressed X-linked genes for the female model were significantly increased and decreased upon Pb exposure, respectively (**Figure 10**). The most upregulated differentially expressed X-linked genes for the female model included *Vma21*, *Nxt2*, *Klhl13*, *Slitrk4*, and *Slitrk2*, while the most downregulated differentially expressed X-linked genes included *Dusp9*, *Abcd1*, *Suv39h1*, *Kdm5c*, and *Stard8* (**Figure 10B** and **Supplementary Figure S13**). The sex-stratified assessment for the male model indicated that 24 (4.49%) of total captured X-linked genes were dysregulated upon Pb exposure (**Table 4** and **Supplementary Table S16**).

**Figure 10.**
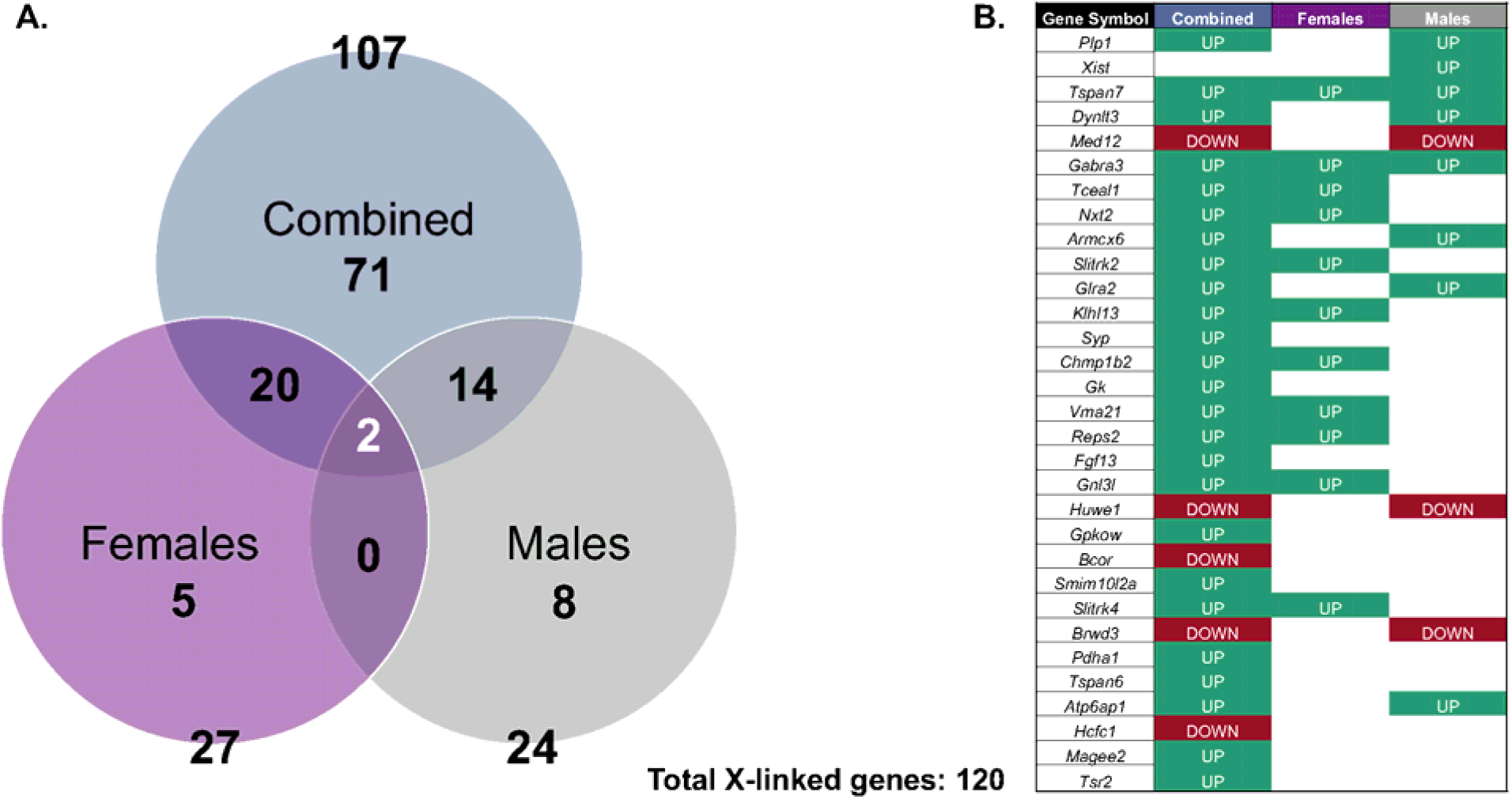
Combined sex, Female, and Male model-specific differentially expressed X-linked gene comparisons. (**A**) Venn diagram of differentially expressed X-linked gene comparisons between each tested model. The combined model shows 107 differentially expressed X-linked genes related to *n*=38 RNA-seq samples, including *n*=18 for control and *n*=20 for the Pb exposure groups. The female/male models show 27/24 differentially expressed X-linked genes, respectively, related to *n*=19 RNA-seq samples, including *n*=9 for control and *n*=10 for the Pb exposure groups. Collectively, 120 differentially expressed X-linked genes were dysregulated by gestational exposure to Pb in the embryonic head. (**B**) The top differentially expressed X-linked genes were recorded based on directionality of gene expression changes. Upregulated (teal; LogFC>0; FDR<0.05) and downregulated (red; LogFC<0; FDR<0.05) differentially expressed X-linked genes upon Pb exposure relative to control are indicated in respective boxes, comparing the three tested models.

Approximately 10 (41.7%) and 14 (58.3%) of the differentially expressed X-linked genes for the male model were significantly increased and decreased upon Pb exposure, respectively (**Figure 10B**). The most upregulated differentially expressed X-linked genes for the male model included *Plp1*, *Xist*, *Dynlt3*, *Tspan7*, and *Armcx6*, while the most downregulated differentially expressed X-linked genes included *Huwe1*, *Med12*, *Aff2*, *Ddx3x*, and *Atrx* (**Figure 10B**), with *Plp1* showing pronounced sexually dimorphic effects of exposure (**Supplementary Figure S14**).

Collectively, 2 (1.7%) of total X-linked genes were common between all three tested models, with 5 (4.2%) and 8 (6.7%) were found exclusively in the female and male models, respectively (**Figure 10A**). A modest but statistically significant positive correlation (r=0.43, *p*<0.001) was observed between the LogFCs of female vs. male models, indicating a tendency for concordance in differentially expressed X-linked gene expression between sexes (**Supplementary Figure S15**). These results suggest that Pb exposure significantly dysregulate X-linked genes by exposure group and sex. The total differentially expressed X-linked genes by exposure group and sex are enriched in GOBP and KEGG pathways, with the heatmap revealing a clear separation by exposure (**Figure 11**). The observed enriched pathways are responsible for metabolism and biosynthetic processes (carbohydrate derivative and organophosphate metabolic processes), gene regulation (negative regulation of transcription by RNA polymerase II and RNA processing), transport and localization (import into cell, inorganic ion transmembrane transport, and membrane organization), development and morphogenesis (chordate embryonic and head development), DNA damage response, and nervous system/diseases (Alzheimer’s disease and synapse organization).

**Figure 11.**
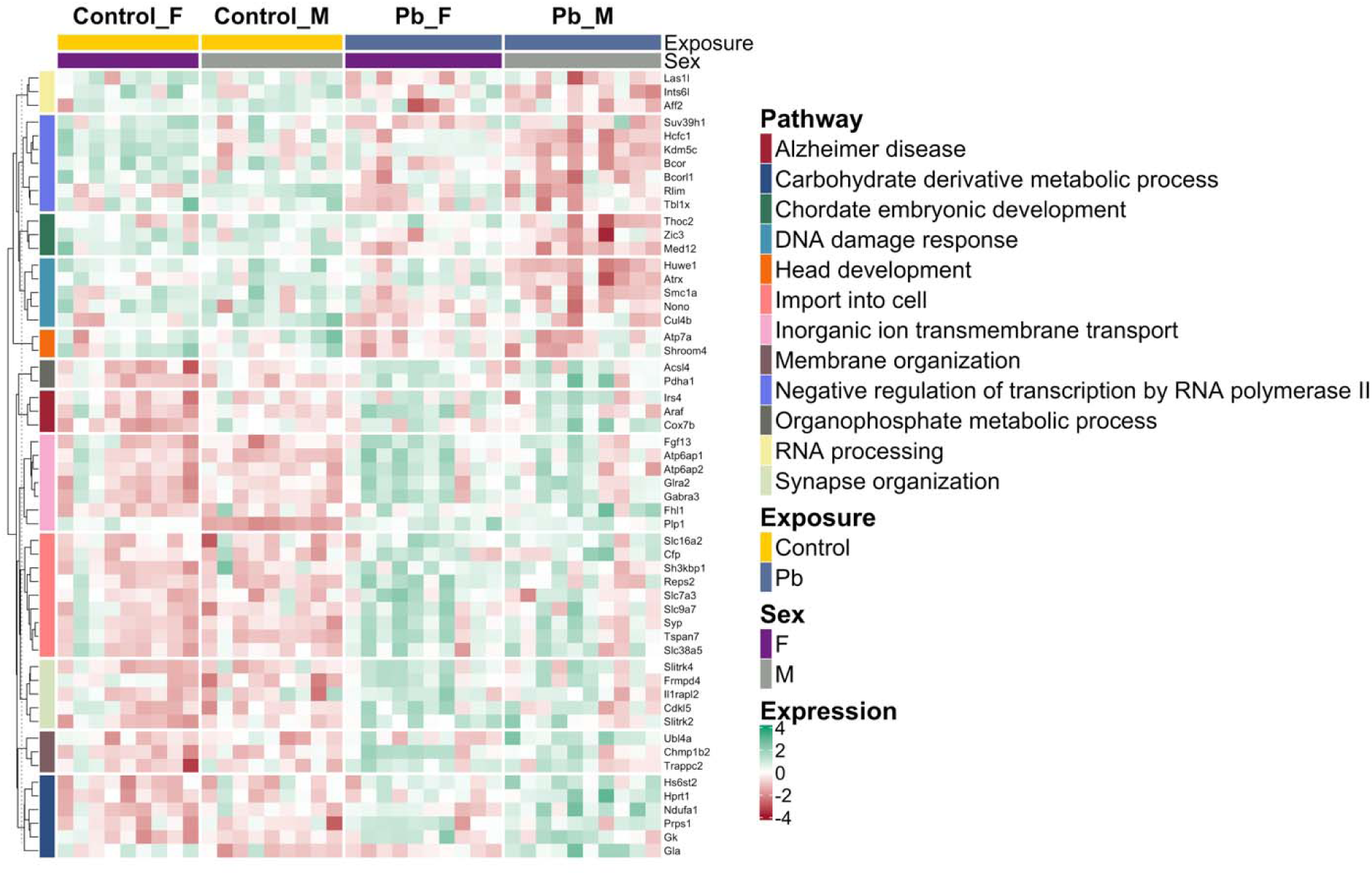
Heatmap of differentially expressed X-linked gene expression within enriched GOBP and KEGG pathways in combined sex and sex-stratified models of Pb exposure. Pathways are indicated by the colored left annotation bar, while exposure (Control: yellow; Pb: dark blue) and sex (Female: F; Male: M) are shown in the top annotation bar. Gene expression levels are represented as z-score scaled log counts per million (logCPM), with gene names displayed on the right. Only GOBP and KEGG pathways containing at least two differentially expressed X-linked genes were included.

### 3.6. Sexually dimorphic autosomal genes are particularly susceptible to gestational Pb exposure in the embryonic head at E13–15

To specifically assess the sexually dimorphic genes within the RNA-seq dataset, differentially expressed genes were identified based on sex exclusively in the control group for *n*=9 females and 9 males. While the control samples cluster distinctly by sex, this pattern disappeared after Pb exposure, as observed by the MDS plots (**Supplementary Figure S16**). For the control sex-specific model, 3.3% (498) of total captured genes were differentially expressed genes in control males relative to control females (**Supplementary Table S17**). Approximately 62.7% (312) and 37.4% (186) of the genes were significantly upregulated and downregulated in control males relative to control females, respectively. The differentially expressed genes identified by sex for the control group indicate sexually dimorphic genes for the embryonic head at E13–15.

The sexually dimorphic autosomal genes were significantly enriched and sensitive to gestational Pb exposure for each sex (*p*=8.8×10^−6^ for females and *p*<2.2×10^−16^ for males; **Table 5**). Our data thus indicates that sexually dimorphic genes are more likely than expected by chance to be transcriptionally responsive to Pb exposure in both sexes. For females, 8.8% (44) of sexually dimorphic genes were significantly dysregulated upon Pb exposure (**Figure 12A**). The most significantly enriched pathways of sexually dimorphic genes dysregulated by Pb exposure for females (**Figure 12B**) include GOCC (CHD-type complex, NuRX complex, and the chromosome, centromeric core domain) and GOMF (GTPase and kinase binding). For males, 15.3% (76) of sexually dimorphic genes were significantly dysregulated upon Pb exposure (**Figure 12C**). The most significantly enriched GOBP pathways of sexually dimorphic genes dysregulated by Pb exposure for males included myelination and axon ensheathment, glial cell development, and cytoskeletal filament organization (**Figure 12D**), while the most significantly enriched KEGG pathways included neurotransmitter and synaptic signaling (amphetamine addiction and cholinergic, dopaminergic, and glutamatergic synapses), Ras signaling pathway, diabetic cardiomyopathy, cellular structure and barrier function (cornfield envelope formation).

**Figure 12.**
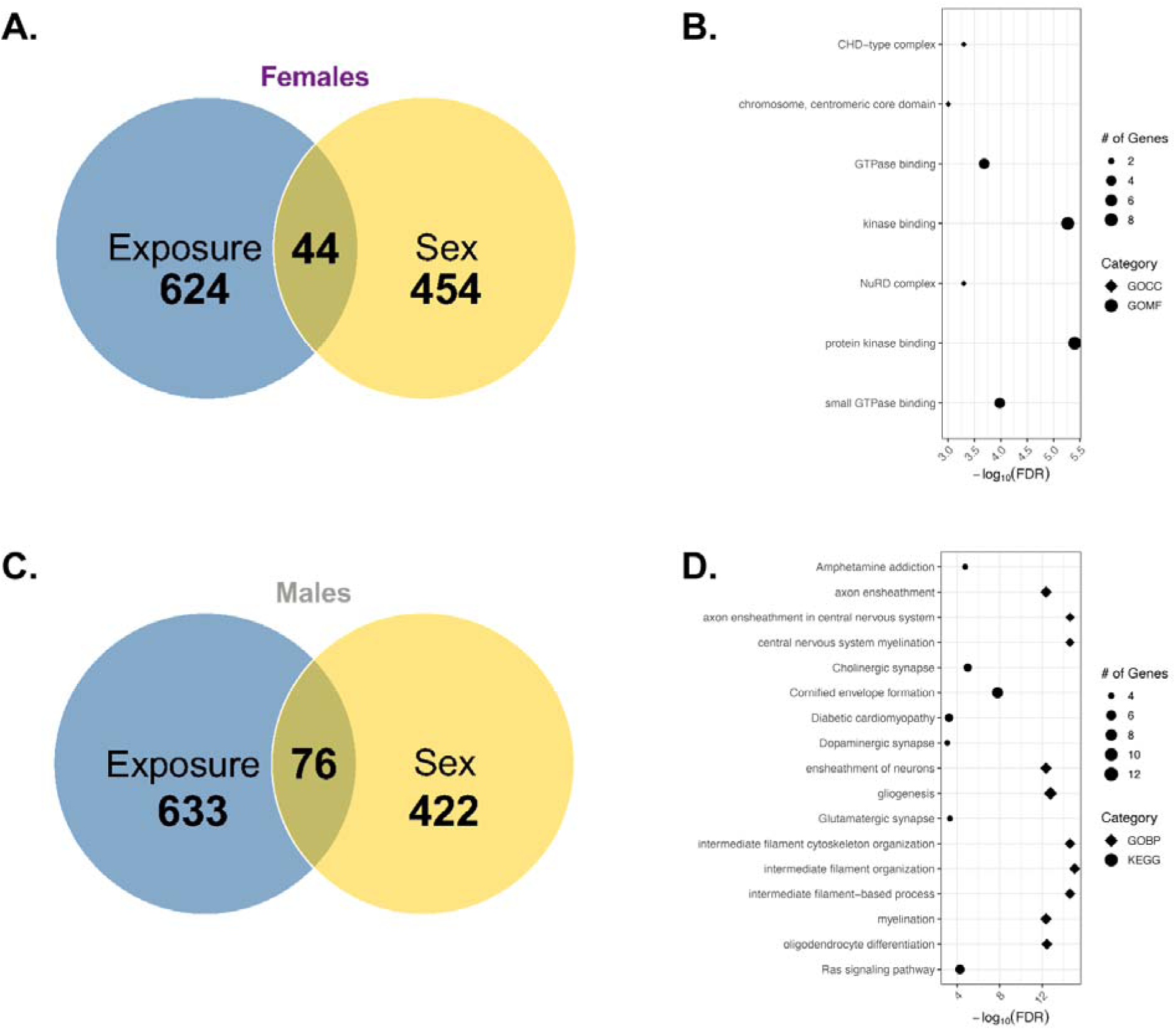
Summary of potential sexually dimorphic genes sensitive to gestational Pb exposure. A total of 498 genes is differentially expressed based on sex at E13–15 in the embryo head data analysis of the control group alone (*n*=9 per sex). According to data presented in the combined model, 668 and 709 genes are differentially expressed based on gestational exposure to Pb at E13–15 in the embryo head in females and males, respectively (*n*=9 per sex for control; *n*=10 per sex for Pb). Data interpretation is based on sex (yellow) and exposure (blue) comparisons for each sex. (**A**) Differentially expressed gene overlap between sex vs. exposure-specific assessment, focusing on females. (**B**) Summary of biological processes significantly dysregulated by Pb exposure for the 44 sexually dimorphic genes, representing the female data. (**C**) Differentially expressed gene overlap between sex vs. exposure-specific assessment, focusing on males. (**D**) Summary of biological processes and KEGG pathways significantly dysregulated by Pb exposure for the 76 sexually dimorphic genes, representing the male data.

**Table 5.**
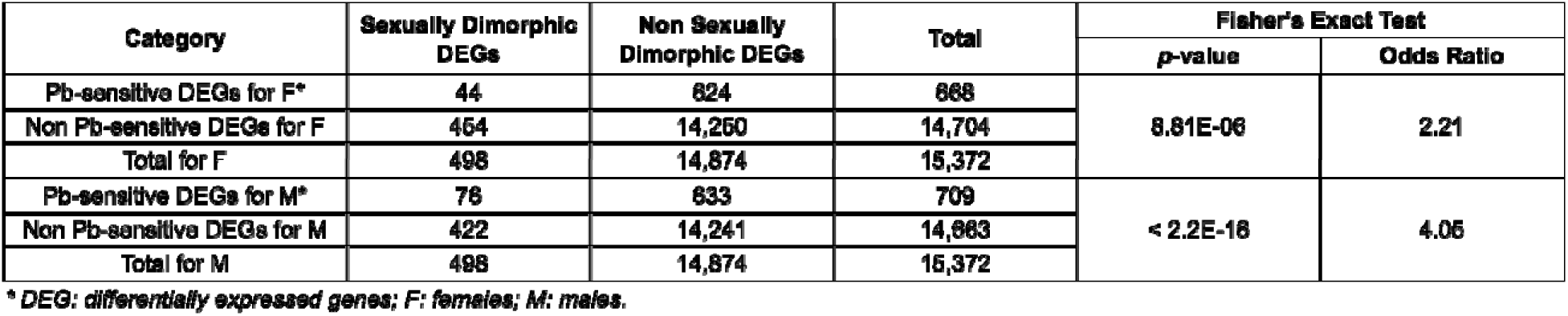
Fisher’s Exact Test for Sexually Dimorphic Differentially Expressed Gene Enrichment for the Embryonic Head at E13–15.

Overall, these data support that preconception and gestational Pb exposure alter neuronal structure, function, and biological processes particularly related to sexually dimorphic genes that may contribute to neurodegenerative diseases.

## 4. DISCUSSION

### 4.1. Key findings

Pb exposure during preconception and gestation significantly increased embryonic weight at E13–15 relative to controls, which was independent of sex and litter. Bulk RNA-sequencing revealed substantial Pb-induced gene expression changes in the mid-gestation embryonic head among all mice and in sex-stratified analyses, including 2,920 differentially expressed genes, 31 affected imprinted genes, and 120 affected X-linked genes that collectively dysregulate neuronal and metabolic pathways and diseases. The current study also reports that sexually dimorphic autosomal genes are particularly sensitive to preconception and gestational Pb exposure, with 44 genes significantly changed in females and 76 changed in males. Pb exposure dysregulated molecular pathways of GOBP (gene expression and RNA metabolism, membrane transport, developmental morphogenesis, and stress responses), GOCC (nuclear structure and gene regulation, synaptic structure and function, and membrane transport complexes), and GOMF (chromatin structure and regulation, neuronal signaling, and protein interactions). The current study also highlights KEGG pathways related to neurotransmission processes, metabolism, cell cycle and cardiometabolic diseases, and neurodegenerative disease risk. These findings suggest that Pb exposure perturbs fetal development by disrupting multiple biological pathways, including those governing imprinting, X-linked genes, and sexually dimorphic autosomal genes. The evidence of such widespread dysregulation underscores the urgent need to elucidate the precise epigenetic regulatory mechanisms involved and to assess the lasting sex-specific impacts on offspring health from birth to adulthood.

### 4.2. The developmental impact of Pb exposure

Our results indicated no significant differences between the number of total embryos, viable embryos, litter sizes, or sex distribution upon Pb exposure at 32 ppm during preconception and gestation. The ratio of males to females were close to an expected Mendelian ratio of 50:50. This observation is consistent with a Swiss mouse model of *in utero* Pb exposure (*n*=6 dams per doses of 14, 28, 56 and 112 mg/kg), which reported that the number of implants, fetuses (alive and dead), and sex ratio were unaffected ^48^. However, embryogenesis is sensitive to Pb, as high doses of 1000–2000 ppm Pb in rats significantly reduces litter sizes and fetal weights ^49^. Our results demonstrate that Pb exposure increases embryonic weight in both sexes (**Table 1** and **Figure 2**). Although this finding is consistent with previous reports of prenatal Pb exposure altering fetal growth parameters, the effects on weight can be heavily dependent on experimental design. For instance, the Swiss mouse study reports no significant differences in fetal body weights at E18, and some epidemiological studies associate maternal blood Pb levels with small-for-gestational-age, low birth weight, and preterm birth ^48,50–53^. As such, these variations in experimental design include differences in the route of exposure, mouse strain, or endpoints measured, as well as the observational nature and inherent limitations of human cohort analyses. Given that our study used a 48-hour, two-night window for mating, it is possible that the observed embryo weight differences arose from Pb-exposed animals mating consistently on the first night and control animals mating consistently on the second night. The likelihood of this scenario is extremely low, as the disparity in embryo weights was observed across n≥9 separate litters per exposure group (**Supplementary** Figure 2). The observed increase in embryonic weight may reflect a compensatory dysregulation in growth or metabolism, aligning with reports that early toxicant exposures trigger adaptive, and potentially maladaptive, responses in the developing embryo ^54^. For a more comprehensive understanding of Pb-induced toxicity and its impact during early development, investigation of maternal-fetal crosstalk is warranted, with a particular focus on the dynamic communication between the maternal brain, the placenta, and the developing embryo ^55^.

### 4.3. Pb exposure dysregulates neuronal gene expression and biological pathways

Pb exposure induced widespread exposure and sex-specific transcriptomic changes in the embryonic head as detected from RNA-seq analysis of all mice and stratified by sex (**Figures 3–5**), collectively showing 2,920 differentially expressed genes (**Table 2** and **Supplementary Figure S6**). The most significantly dysregulated differentially expressed genes by exposure group (**Figure 3**) were not sex dependent. Among these, *Gsdme* is a tumor suppressor gene involved in pyroptosis and inflammatory responses, and its upregulation suggests a generalized stress or cell death pathways activated by Pb in the developing brain ^56^. Conversely, *Supt16*, a chromatin remodeler, was the most significantly downregulated differentially expressed gene observed, suggesting broad repressive effects on transcriptional regulation and neurodevelopment associated with Pb exposure ^57,58^. The biological processes associated with Pb exposure (**Figure 7**) suggest a shift towards neuronal synaptic activity and reduced gene regulatory plasticity during development. For instance, the upregulation of synaptic signaling and vesicle-mediated transport, neurotransmitter transport, and ion transmembrane transport points to an increased demand for efficient synaptic communication, while the downregulation of mRNA metabolic processes, chromatin organization, and embryonic development pathways indicates suppression of neuronal plasticity at a transcriptional and epigenetic level. According to the KEGG results, these transcriptional changes are associated with neurodegenerative and addiction disease processes (including Parkinson’s disease and nicotine addiction). The enriched pathways identified in our study is consistent with known mechanisms of Pb neurotoxicity and disease associations. Pb disrupts presynaptic vesicular release by altering synaptic proteins, inhibits N-methyl-D-aspartate receptor (NMDAR) signaling, and impairs brain-derived neurotrophic factor (BDNF) release, thereby reducing synaptic plasticity and function ^59^. There is evidence confirming Pb exposure as a significant risk factor for Parkinson’s disease ^60^. Thus, our results linking synaptic hyperactivity and oxidative metabolism upregulation with suppressed gene regulatory systems echo these toxicological insights and highlight a mechanistic basis for Pb-related cognitive decline.

In the female model (**Figure 4**), dysregulated *Scn2b* (encoding a subunit of the voltage-gated sodium channel), *Wnt4* (involved in key developmental signaling pathways and homeostasis in the brain), and *Trpc2* (a cation channel linked to sex-specific social behaviors), point to potential impairments in neuronal signaling pathways critical for development and sensory processing ^61–63^. Collectively, the female-specific gene expression changes are involved in overall downregulation of DNA metabolic processes, contributing to genome instability and cellular dysfunction, while upregulation of glutamatergic synapse-related disease pathways suggest disruption or heightened neuronal excitatory signaling (**Figure 7**).

The most notable trends were observed in the male model (**Figure 5**), indicating upregulation of myelin-related genes, including *Mobp* (myelin-associated oligodendrocytic basic protein), *Mbp* (myelin basic protein), and *Plp1* (Proteolipid protein 1; an essential component of myelin). Excessive levels of these proteins underlie several neurodegenerative conditions and psychiatric disorders ^64,65^. Our findings closely align with a reported single-cell RNA-seq analysis indicating that perinatal Pb exposure is associated with a 12.4% increase in oligodendrocyte abundance, even following a four-month cessation of 32 ppm Pb exposure in mice ^66^. Interestingly, these oligodendrocyte markers were also sensitive to gestational nicotine exposure in rats in a sex-specific manner ^67^. On the other hand, striking differentially expressed gene patterns of male downregulation were dominated by keratin genes (*Krt6b*, *Krt81*, *Krt83*, *Krt6a*), which can serve as stress response proteins, and this downregulation might reduce protection against Pb-induced damage in epithelial tissues ^68^. Our pathway analysis for males revealed distinct biological disruptions associated with Pb exposure, reflecting both DNA damage responses and altered metabolic processes (**Figure 7**). Despite the lack of a sex-specific assessment of Pb-induced DNA damage response, our data is consistent with previous literature showing that Pb impairs DNA repair mechanisms by downregulating key repair genes involved in base excision repair, nucleotide excision repair, and double-strand break repair, thus contributing to mutagenic outcomes ^69^. The male-specific KEGG pathway upregulation of the TCA cycle suggests a shift in cellular metabolism, likely due to increased energy demands under Pb-induced oxidative stress, while the downregulation of PRC pathways suggests that Pb-induced epigenetic dysregulation of chromatin remodelers is likely involved in transcriptional repression.

Taken together, our results suggest that Pb exposure in females may induce greater disruption in regulatory neuronal signaling and developmental control, while males may undergo more neurostructural and cytoskeletal alterations. Certain gene expression changes (such as upregulated *Gh*) could plausibly support enhanced growth and subsequent weight gain in Pb-exposed samples. However, the simultaneous dysregulation of multiple genes essential to development and tissue homeostasis identified in enriched pathways (**Figure 6**) makes it difficult to pinpoint the basis of Pb neurotoxicity. Future studies integrating longitudinal transcriptional profiles with functional assays of synaptic and oxidative stress will be essential to define causality and identify therapeutic targets for later-life neurological diseases.

### 4.4. Pb exposure disrupts imprinted genes and X-linked genes

Even though imprinted genes and X-linked genes represent a subset of the transcriptome, they play crucial roles for proper embryonic development. Pb-induced exposure and sex-specific differentially expressed genes are associated with 31 imprinted genes (**Table 3**, **Figure 8**, and **Supplementary Figures S8–S11**). Imprinted genes cluster at imprinted domains that typically cover 100–1700 kb genomic regions, including at least one lncRNA, differentially methylated regions, and an imprinting control region (ICR) that regulates the entire canonical imprinting domains ^8^. Thus, each imprinting cluster is regulated through distinct epigenetic mechanisms such as the lncRNA-mediated silencing (insulin-like growth factor 2 receptor; *Airn*/*Igf2r* cluster) and the insulator (H19/*Igf2* cluster) models. ICRs have been shown to be sensitive to Pb exposure during development in human and cell culture models ^20–22,25^. Our results provide strong evidence that imprinted genes are dysregulated following gestational Pb exposure in the developing embryonic brain that include maternally and paternally imprinted genes (**Figure 8**). The Angelman syndrome and Prader-Willi syndrome imprinted domain cluster is in the same genomic region, where the antisense RNA (*Ube3a*-ATS) originates from the paternal *Snrpn* promoter involved in the imprinting setting of the *Ube3a* paternal allele in the brain ^70^. As indicated by the differential expression of *Ube3a* (downregulated in males) and *Snrpn* (upregulated in combined sex) and their associated pathways in sexual reproduction (**Figures 8** and **9**), Pb-induced transcriptional dysregulation may impact this mechanism, but confirmation of a direct association is needed. Along with *de novo* DNAm, ncRNA including PIWI-interacting RNA (piRNA) and histone modifications can also contribute to non-canonical imprinting regulation ^71^. The upstream genomic regions of the Ras protein-specific guanine nucleotide releasing factor 1 (*Rasgrf1*) generate piRNA that recruits DNAm at the *Rasgrf1*-ICR to maintain its paternal imprinting status ^72^. In this study, *Rasgrf1* is among the most significantly upregulated imprinted genes following gestational Pb exposure (**Figure 8**), with its altered expression linked to disrupted synaptic signaling pathways in the embryonic brain (**Figure 9**). Our previous work demonstrated that piRNA is highly expressed in the hippocampus and that perinatal Pb exposure (32 ppm) alters piRNA expression in the male brain, suggesting that Pb influences piRNA-mediated epigenetic regulation ^26,73^. Similarly, perinatal Pb exposure at the same dose and time window alters heart-specific piRNA expression in a sex-specific manner, further suggesting that Pb-induced piRNA dysregulation during development may influence cardiovascular disease risk ^34^. These findings, together with existing literature, highlight the critical role of *Rasgrf1* in hippocampal memory formation and its calcium-dependent activation via NMDAR signaling ^74^. The impact of developmental Pb exposure on imprinted genes and the consequences for sex differences in neurotoxicity remain largely unexplored and warrant further investigation.

Sex linked genes perform essential roles in sexual differentiation and drive sex-specific brain development even before gonadal formation ^75^. In this study, Pb exposure and sex-specific differentially expressed genes were associated with 120 X-linked genes (**Table 4**, **Figure 10**, and **Supplementary Figures S12–S15**). Notably, **Figure 11** highlights the overwhelming impact of Pb toxicity on these genes and illustrates that the most significantly dysregulated X-linked genes are linked to key biological and disease pathways. The oligodendrocyte marker *Plp1* is among the most differentially expressed X-linked gene following Pb exposure, and is associated with inorganic ion transmembrane transport, suggesting its role in neuronal signaling and overall neuronal cellular functions. One of the most enriched KEGG pathways linked in this analysis is Alzheimer’s disease (**Figure 11**), suggesting that gestational Pb exposure may contribute to early molecular changes associated with neurodegenerative risk and underscoring the long-term implications of X-linked gene dysregulation by Pb. This finding is consistent with some research suggesting that early life Pb exposure is linked to Alzheimer’s disease risk ^5,76^. X-linked imprinted genes contribute to sex differences in brain function and may increase vulnerability to disease when affected by Pb exposure ^23^. Normally, *Xist* is a prominent X-linked imprinted gene exclusively expressed in females and involved in random XCI; it is often used as a marker for sex genotyping. Our results are consistent with this, showing high *Xist* expression in females with no significant difference between control and Pb-exposed groups (**Figure 10** and **Supplementary Figure S14**). In contrast, *Xist* expression in males is nearly absent under control conditions but increases sharply (by a 2.9 log fold change) following Pb exposure (**Supplementary Table S16**). Further research is needed to determine whether this transcriptional change results from Pb-induced de-repression via altered DNAm or disrupted histone modifications. Given the complex epigenetic mechanisms involved in the process of genomic imprinting, interpretation of our findings on Pb-induced changes in imprinted and X-linked gene expression should be approached with caution and incorporated in future studies, as these results must consider the intricate nature of imprinting clusters, sex-, tissue-, developmental stage-specific regulation, and Pb-induced neurotoxicity.

### 4.5. Pb exposure elicits sexually dimorphic responses

Differential gene expression between the sexes, including sex-linked and autosomal genes, are influenced by the unique content of the sex chromosomes ^31^. In the exposure-specific comparison of differentially expressed genes (control vs. Pb-exposed groups), we observed a sex-stratified impact on Pb toxicity (**Supplementary Figure S5**), suggesting that sexually dimorphic autosomal genes are particularly sensitive to Pb exposure. This observation was further supported by the sex-specific comparison of differentially expressed genes using the same dataset (control females vs. male groups; **Supplementary Figure S16**). In the control group, samples cluster by sex, indicating underlying biological or molecular distinctions between males and females. The suppression of this trend by Pb exposure suggests that it either masks, reduces, or overrides these inherent sex-based differences. The current study reports that sexually dimorphic genes are particularly sensitive to preconception and gestational Pb exposure, males appear to be more susceptible to Pb-induced changes in sexually dimorphic gene expression, with 76 genes significantly altered in males compared to 44 in females (**Figure 12**).

Collectively, our findings reveal that Pb-induced sexually dimorphic gene expression changes are linked to distinct molecular pathways including chromatin remodeling processing in females; changes in males were associated with neural structure and functional processes, as well as those associated with nervous system and cardiovascular diseases (**Figure 12**). The sexually dimorphic gene responses observed in our study echo reports demonstrating that males and females differ in their transcriptomic and phenotypic susceptibility to Pb ^22,23,30^. Animal studies often indicate greater or unique neurobehavioral, metabolic, or epigenetic impacts in one sex following gestational Pb exposure ^29,30,77^. For example, one study found that gestational Pb exposure is associated to male-specific late-onset obesity and decreased spontaneous motor activity, implicating cholinergic and glutamatergic neurons (cell populations highlighted in male-specific assessment in our study) as being affected by Pb exposure ^29^. Another study demonstrates that molecular and neurobehavioral effects of gestational Pb exposure is parent and sex-dependent, further suggesting that Pb-induced sexually dimorphic effects are likely regulated via epigenetic mechanisms ^77^. There is extensive research indicating Pb-induced sexually dimorphic effects in cardiovascular disease following gestational exposure ^22,30,78^. Taken together, these insights emphasize the critical need to consider sex as a biological variable in assessing the neurodevelopmental and metabolic risks of early life Pb exposure, as targeted interventions may be necessary to protect the most vulnerable populations.

## 5. STRENGTHS AND LIMITATIONS

The current study is among the first to identify significant association between perinatal Pb exposure effects, including increased embryonic weight at E13–15 and expression changes in sexually dimorphic genes in the embryonic head, specifically at a human exposure-relevant dose. By carefully cataloging the whole embryonic weight phenotypes using linear regression models and subsequent RNA-seq assessment, this study explored genome-wide gene expression differences upon Pb exposure (in the combined sex model) and sex (male and female models) during a critical exposure time window at murine mid-gestation. Findings from this study fill a critical knowledge gap in understanding the association between Pb toxicity and gestational neurodevelopment as it relates to genomic imprinting, sex-specific effects on gene expression in the brain and the associated biological and disease pathways.

There are several limitations to the current study. First, while this study was designed using the embryonic head as a proxy to capture gene expression changes in the developing brain, it may also contain other tissues such as the connective tissues, craniofacial skeleton, and mesenchyme and connective tissues among others, that could contribute to the obtained results. Second, although bulk-RNA seq captured genome-wide gene expression, the method is unable to detect cell type-specific changes linked to Pb exposure. Thus, future work will address how these gene expression alterations perturb temporal neurodevelopmental functions. Third, the current study is conducted using a murine model from the C57BL/6J wildtype background, which does not differentiate between the allele-specific gene expression changes. As such, future confirmation is needed for the allelic contribution of gene expression changes linked to Pb toxicity, especially for imprinted gene assessment. Fourth and finally, as the study design draws conclusions of Pb-induced increased embryonic weight and its association to sexually dimorphic gene expression changes and pathways, the establishment of causality is limited. As such, the associations should be interpreted cautiously. Future research is encouraged to confirm the overall epigenetic pathways of Pb-induced toxicity, including the evaluation of DNAm, histone modifications, and ncRNA in the maternal-fetal crosstalk. Similarly, repeated measurements of Pb-associated gene expression changes in the brain using mouse models of disease relevance at multiple timepoints in development would provide a robust causal inference to the disease pathways highlighted in this study.

## 6. CONCLUSION

This study demonstrates that Pb exposure during preconception and gestation, corresponding to a human-relevant maternal blood Pb level average of 9.7 μg/dL, disrupts key developmental processes by broadly changing embryonic gene expression, including imprinted genes, X-linked genes, and neuronal and metabolic pathways. Taken together, these molecular alterations and pathways are linked to the Pb-induced increased embryonic weight phenotype. Furthermore, these molecular disturbances implicate Pb as a potent developmental toxicant, affecting pathways fundamental to brain function, metabolism, and overall embryonic development. The pronounced sensitivity of sexually dimorphic genes to gestational Pb exposure highlights potential sex-specific vulnerabilities and associated long-term health impacts of early life exposure. As such, a deeper investigation into Pb toxicity and its effects on epigenetic programming will lay the foundation for developing targeted interventions to alleviate long-term health consequences of early life Pb exposure. Future research will assess epigenetic mechanisms altered by Pb exposure to inform strategies for alleviating its associated adverse health outcomes instigated during development.

## Authorship contribution statement

BPUP, DCD, and JAC conceived and designed the experiments. BPUP, AT, JP, MAS, DCD, JAC planned the experimental setup, while BPUP, AT, JP, DW, TG, RKM, and JAC carried out various aspects of the experiments. BPUP, ML, AT, JP, DW, and JAC contributed to experimental data collection and curation. BPUP, ML, TG, JMG, and JAC analyzed and interpreted the results. BPUP, KMB, JMG, MAS, DCD, and JAC supervised the experimental work. BPUP, ML, AT, and JAC drafted the original manuscript, with JP, DW, TG, RKM, KMB, JMG, MAS, and DCD contributing to writing and revision of the manuscript. All authors have read and approved the final version of the manuscript draft.

## Acknowledgements

The authors would like to thank Dr. Sundeep Kalantry and Dr. Vasantha Padmanabhan for their thoughtful discussion of results interpretation, the University of Michigan Advance Genomics Core for their assistance with RNA sequencing, and the University of Michigan Unit for Laboratory Animal Medicine for ensuring proper animal care. We would like to also thank Dr. Mathia Colwell and Jennifer Smith for their assistance during animal dissections.

## Data availability

The data submission for GEO and Github are in progress.

## REFERENCES

1. Bhasin T, Lamture Y, Kumar M, Dhamecha R. Unveiling the Health Ramifications of Lead Poisoning: A Narrative Review. Cureus. Oct 2023;15(10):e46727. doi:10.7759/cureus.46727

2. Chowdhury R, Ramond A, O’Keeffe LM, et al. Environmental toxic metal contaminants and risk of cardiovascular disease: systematic review and meta-analysis. BMJ. Aug 29 2018;362:k3310. doi:10.1136/bmj.k3310

3. Ruckart PZ, Jones RL, Courtney JG, et al. Update of the Blood Lead Reference Value – United States, 2021. MMWR Morb Mortal Wkly Rep. Oct 29 2021;70(43):1509–1512. doi:10.15585/mmwr.mm7043a4

4. Tung PW, Burt A, Karagas M, et al. Association between placental toxic metal exposure and NICU Network Neurobehavioral Scales (NNNS) profiles in the Rhode Island Child Health Study (RICHS). Environ Res. 03 2022;204(Pt A):111939. doi:10.1016/j.envres.2021.111939

5. Santa Maria MP, Hill BD, Kline J. Lead (Pb) neurotoxicology and cognition. Appl Neuropsychol Child. 2019 Jul-Sep 2019;8(3):272–293. doi:10.1080/21622965.2018.1428803

6. Nishioka E, Yokoyama K, Matsukawa T, et al. Evidence that birth weight is decreased by maternal lead levels below 5μg/dl in male newborns. Reprod Toxicol. Aug 2014;47:21–6. doi:10.1016/j.reprotox.2014.05.007

7. Robles-Matos N, Artis T, Simmons RA, Bartolomei MS. Environmental Exposure to Endocrine Disrupting Chemicals Influences Genomic Imprinting, Growth, and Metabolism. Genes (Basel). 07 28 2021;12(8)doi:10.3390/genes12081153

8. Barlow DP, Bartolomei MS. Genomic imprinting in mammals. Cold Spring Harb Perspect Biol. Feb 2014;6(2)doi:10.1101/cshperspect.a018382

9. Schaafsma SM, Pfaff DW. Etiologies underlying sex differences in Autism Spectrum Disorders. Frontiers in neuroendocrinology. Aug 2014;35(3):255–71. doi:10.1016/j.yfrne.2014.03.006

10. Engel E. Uniparental disomy (UPD). Genomic imprinting and a case for new genetics (prenatal and clinical implications: the “Likon” concept). Annales de genetique. 1997;40(1):24–34.

11. Monk D, Mackay DJG, Eggermann T, Maher ER, Riccio A. Genomic imprinting disorders: lessons on how genome, epigenome and environment interact. Nature reviews Genetics. 04 2019;20(4):235–248. doi:10.1038/s41576-018-0092-0

12. Goovaerts T, Steyaert S, Vandenbussche CA, et al. A comprehensive overview of genomic imprinting in breast and its deregulation in cancer. Nat Commun. 10 2018;9(1):4120. doi:10.1038/s41467-018-06566-7

13. Rabinovitz S, Kaufman Y, Ludwig G, Razin A, Shemer R. Mechanisms of activation of the paternally expressed genes by the Prader-Willi imprinting center in the Prader-Willi/Angelman syndromes domains. Proceedings of the National Academy of Sciences of the United States of America. May 8 2012;109(19):7403–8. doi:10.1073/pnas.1116661109

14. Cassidy SB, Schwartz S, Miller JL, Driscoll DJ. Prader-Willi syndrome. Genet Med. Jan 2012;14(1):10–26. doi:10.1038/gim.0b013e31822bead0

15. Margolis SS, Sell GL, Zbinden MA, Bird LM. Angelman Syndrome. Neurotherapeutics. Jul 2015;12(3):641–50. doi:10.1007/s13311-015-0361-y

16. Cassidy FC, Charalambous M. Genomic imprinting, growth and maternal-fetal interactions. J Exp Biol. 03 07 2018;221(Pt Suppl 1)doi:10.1242/jeb.164517

17. Hudson QJ, Kulinski TM, Huetter SP, Barlow DP. Genomic imprinting mechanisms in embryonic and extraembryonic mouse tissues. Heredity (Edinb). Jul 2010;105(1):45–56. doi:10.1038/hdy.2010.23

18. Perera BP, Teruyama R, Kim J. Yy1 gene dosage effect and bi-allelic expression of Peg3. PLoS One. 2015;10(3):e0119493. doi:10.1371/journal.pone.0119493

19. Kim J, Frey WD, He H, et al. Peg3 mutational effects on reproduction and placenta-specific gene families. PLoS One. 2013;8(12):e83359. doi:10.1371/journal.pone.0083359

20. Goodrich JM, Sánchez BN, Dolinoy DC, et al. Quality control and statistical modeling for environmental epigenetics: a study on in utero lead exposure and DNA methylation at birth. Epigenetics. 2015;10(1):19–30. doi:10.4161/15592294.2014.989077

21. Nye MD, King KE, Darrah TH, et al. Maternal blood lead concentrations, DNA methylation of. Environ Epigenet. 2016;2(1)doi:10.1093/eep/dvv009

22. Svoboda LK, Neier K, Wang K, et al. Tissue and sex-specific programming of DNA methylation by perinatal lead exposure: implications for environmental epigenetics studies. Epigenetics. Oct 2021;16(10):1102–1122. doi:10.1080/15592294.2020.1841872

23. Singh G, Singh V, Sobolewski M, Cory-Slechta DA, Schneider JS. Sex-Dependent Effects of Developmental Lead Exposure on the Brain. Front Genet. 2018;9:89. doi:10.3389/fgene.2018.00089

24. Morgan RK, Tapaswi A, Polemi KM, et al. Environmentally Relevant Lead Exposure Impacts Gene Expression in SH-SY5Y Cells Throughout Neuronal Differentiation. Toxicol Sci. May 21 2025;doi:10.1093/toxsci/kfaf072

25. Morgan RK, Wang K, Svoboda LK, et al. Effects of Developmental Lead and Phthalate Exposures on DNA Methylation in Adult Mouse Blood, Brain, and Liver: A Focus on Genomic Imprinting by Tissue and Sex. Environ Health Perspect. Jun 2024;132(6):67003. doi:10.1289/EHP14074

26. Perera BPU, Wang K, Wang D, et al. Sex and tissue-specificity of piRNA regulation in adult mice following perinatal lead (Pb) exposure. Epigenetics. Dec 2024;19(1):2426952. doi:10.1080/15592294.2024.2426952

27. Raznahan A, Disteche CM. X-chromosome regulation and sex differences in brain anatomy. Neurosci Biobehav Rev. Jan 2021;120:28–47. doi:10.1016/j.neubiorev.2020.10.024

28. Malcore RM, Kalantry S. A Comparative Analysis of Mouse Imprinted and Random X-Chromosome Inactivation. Epigenomes. Feb 10 2024;8(1)doi:10.3390/epigenomes8010008

29. Leasure JL, Giddabasappa A, Chaney S, et al. Low-level human equivalent gestational lead exposure produces sex-specific motor and coordination abnormalities and late-onset obesity in year-old mice. Environ Health Perspect. Mar 2008;116(3):355–61. doi:10.1289/ehp.10862

30. Svoboda LK, Wang K, Goodrich JM, et al. Perinatal Lead Exposure Promotes Sex-Specific Epigenetic Programming of Disease-Relevant Pathways in Mouse Heart. Toxics. Jan 16 2023;11(1)doi:10.3390/toxics11010085

31. Grath S, Parsch J. Sex-Biased Gene Expression. Annual review of genetics. Nov 23 2016;50:29–44. doi:10.1146/annurev-genet-120215-035429

32. Kasten-Jolly J, Lawrence DA. Sex-specific effects of developmental lead exposure on the immune-neuroendocrine network. Toxicol Appl Pharmacol. Nov 1 2017;334:142–157. doi:10.1016/j.taap.2017.09.009

33. Svoboda LK, Perera BPU, Morgan RK, Polemi KM, Pan J, Dolinoy DC. Toxicoepigenetics and Environmental Health: Challenges and Opportunities. Chem Res Toxicol. Aug 15 2022;35(8):1293–1311. doi:10.1021/acs.chemrestox.1c00445

34. Sala-Hamrick KE, Wang K, Perera BPU, Sartor MA, Svoboda LK, Dolinoy DC. Sex-stratified piRNA expression analysis reveals shared functional impacts of perinatal lead (Pb) exposure in murine hearts. Epigenetics. Dec 2025;20(1):2542879. doi:10.1080/15592294.2025.2542879

35. Ward JM, Elmore SA, Foley JF. Pathology methods for the evaluation of embryonic and perinatal developmental defects and lethality in genetically engineered mice. Vet Pathol. Jan 2012;49(1):71–84. doi:10.1177/0300985811429811

36. Yamamoto S, Nagao Y, Kuroiwa K, et al. Rapid selection of XO embryonic stem cells using Y chromosome-linked GFP transgenic mice. Transgenic Res. Oct 2014;23(5):757–65. doi:10.1007/s11248-014-9813-0

37. Alexandra Kuznetsova PBB, Rune H. B. Christensen. lmerTest Package: Tests in Linear Mixed Effects Models. Journal of Statistical Software. 2017;82(13):1–26. doi:doi: 10.18637/jss.v082.i13

38. Douglas Bates MM, Benjamin M. Bolker, Steven C. Walker. Fitting Linear Mixed-Effects Models Using lme4. Journal of Statistical Software. 2015;67(1):1–48. doi:doi:10.18637/jss.v067.i01

39. Sala-Hamrick KE, Tapaswi A, Polemi KM, Nguyen VK, Colacino JA. High-Throughput Transcriptomics of Nontumorigenic Breast Cells Exposed to Environmentally Relevant Chemicals. Environ Health Perspect. Apr 2024;132(4):47002. doi:10.1289/EHP12886

40. Andrews S. FASTQC. A quality control tool for high throughput sequence data. 2010.

41. Love MI, Soneson C, Patro R. Swimming downstream: statistical analysis of differential transcript usage following Salmon quantification. F1000Res. 2018;7:952. doi:10.12688/f1000research.15398.3

42. Robinson MD, McCarthy DJ, Smyth GK. edgeR: a Bioconductor package for differential expression analysis of digital gene expression data. Bioinformatics. Jan 1 2010;26(1):139–40. doi:btp616 [pii] 10.1093/bioinformatics/btp616

43. Ritchie ME, Phipson B, Wu D, et al. limma powers differential expression analyses for RNA-sequencing and microarray studies. Nucleic Acids Res. Apr 20 2015;43(7):e47. doi:10.1093/nar/gkv007

44. Sartor MA, Leikauf GD, Medvedovic M. LRpath: a logistic regression approach for identifying enriched biological groups in gene expression data. Bioinformatics. Jan 15 2009;25(2):211–7. doi:10.1093/bioinformatics/btn592

45. Gu Z, Eils R, Schlesner M. Complex heatmaps reveal patterns and correlations in multidimensional genomic data. Bioinformatics. Sep 15 2016;32(18):2847–9. doi:10.1093/bioinformatics/btw313

46. Yu G, Wang LG, Han Y, He QY. clusterProfiler: an R package for comparing biological themes among gene clusters. OMICS. May 2012;16(5):284–7. doi:10.1089/omi.2011.0118

47. McFarland MJ, Hauer ME, Reuben A. Half of US population exposed to adverse lead levels in early childhood. Proceedings of the National Academy of Sciences of the United States of America. Mar 15 2022;119(11):e2118631119. doi:10.1073/pnas.2118631119

48. Fuentes M, Torregrosa A, Mora R, Gotzens V, Corbellla J, Domingo JL. Placental effects of lead in mice. Placenta. Jul-Aug 1996;17(5-6):371–6. doi:10.1016/s0143-4004(96)90063-6

49. Singh C, Saxena DK, Murthy RC, Chandra SV. Embryo-fetal development influenced by lead exposure in iron-deficient rats. Hum Exp Toxicol. Jan 1993;12(1):25–8. doi:10.1177/096032719301200105

50. Rodosthenous RS, Burris HH, Svensson K, et al. Prenatal lead exposure and fetal growth: Smaller infants have heightened susceptibility. Environ Int. Feb 2017;99:228–233. doi:10.1016/j.envint.2016.11.023

51. Vigeh M, Sahebi L, Yokoyama K. Prenatal blood lead levels and Birth Weight: a Meta-analysis study. J Environ Health Sci Eng. Jun 2023;21(1):1–10. doi:10.1007/s40201-022-00843-w

52. Cheng L, Zhang B, Huo W, et al. Fetal exposure to lead during pregnancy and the risk of preterm and early-term deliveries. Int J Hyg Environ Health. Aug 2017;220(6):984–989. doi:10.1016/j.ijheh.2017.05.006

53. Vigeh M, Yokoyama K, Seyedaghamiri Z, et al. Blood lead at currently acceptable levels may cause preterm labour. Occup Environ Med. Mar 2011;68(3):231–4. doi:10.1136/oem.2009.050419

54. Gluckman PD, Hanson MA, Cooper C, Thornburg KL. Effect of in utero and early-life conditions on adult health and disease. N Engl J Med. Jul 3 2008;359(1):61–73. doi:10.1056/NEJMra0708473

55. Keverne EB. Genomic imprinting, action, and interaction of maternal and fetal genomes. Proceedings of the National Academy of Sciences of the United States of America. Jun 2 2015;112(22):6834–40. doi:10.1073/pnas.1411253111

56. Jiang M, Qi L, Li L, Li Y. The caspase-3/GSDME signal pathway as a switch between apoptosis and pyroptosis in cancer. Cell Death Discov. 2020;6:112. doi:10.1038/s41420-020-00349-0

57. Mylonas C, Tessarz P. Transcriptional repression by FACT is linked to regulation of chromatin accessibility at the promoter of ES cells. Life Sci Alliance. Jun 2018;1(3):e201800085. doi:10.26508/lsa.201800085

58. Wang J, Zhu X, Dai L, et al. Supt16 haploinsufficiency causes neurodevelopment disorder by disrupting MAPK pathway in neural stem cells. Human molecular genetics. Feb 19 2023;32(5):860–872. doi:10.1093/hmg/ddac240

59. Neal AP, Stansfield KH, Worley PF, Thompson RE, Guilarte TR. Lead exposure during synaptogenesis alters vesicular proteins and impairs vesicular release: potential role of NMDA receptor-dependent BDNF signaling. Toxicol Sci. Jul 2010;116(1):249–63. doi:10.1093/toxsci/kfq111

60. Shvachiy L, Geraldes V, Outeiro TF. Uncovering the Molecular Link Between Lead Toxicity and Parkinson’s Disease. Antioxid Redox Signal. Aug 2023;39(4-6):321–335. doi:10.1089/ars.2022.0076

61. Shimada Y, Sato T, Yajima T, et al. SCN2B in the Rat Trigeminal Ganglion and Trigeminal Sensory Nuclei. Cell Mol Neurobiol. Nov 2016;36(8):1399–1408. doi:10.1007/s10571-016-0340-9

62. Liu J, Xiao Q, Xiao J, et al. Wnt/beta-catenin signalling: function, biological mechanisms, and therapeutic opportunities. Signal Transduct Target Ther. Jan 3 2022;7(1):3. doi:10.1038/s41392-021-00762-6

63. Pfau DR, Baribeau S, Brown F, Khetarpal N, Marc Breedlove S, Jordan CL. Loss of TRPC2 function in mice alters sex differences in brain regions regulating social behaviors. J Comp Neurol. Oct 2023;531(15):1550–1561. doi:10.1002/cne.25528

64. Wasinski F, Tavares MR, Gusmao DO, et al. Central growth hormone action regulates neuroglial and proinflammatory markers in the hypothalamus of male mice. Neurosci Lett. May 29 2023;806:137236. doi:10.1016/j.neulet.2023.137236

65. Rajkowska G, Mahajan G, Maciag D, et al. Oligodendrocyte morphometry and expression of myelin – Related mRNA in ventral prefrontal white matter in major depressive disorder. J Psychiatr Res. Jun 2015;65:53–62. doi:10.1016/j.jpsychires.2015.04.010

66. Bakulski KM, Dou JF, Thompson RC, et al. Single-Cell Analysis of the Gene Expression Effects of Developmental Lead (Pb) Exposure on the Mouse Hippocampus. Toxicol Sci. 08 2020;176(2):396–409. doi:10.1093/toxsci/kfaa069

67. Cao J, Wang J, Dwyer JB, et al. Gestational nicotine exposure modifies myelin gene expression in the brains of adolescent rats with sex differences. Transl Psychiatry. Apr 16 2013;3(4):e247. doi:10.1038/tp.2013.21

68. Kalabusheva EP, Shtompel AS, Rippa AL, Ulianov SV, Razin SV, Vorotelyak EA. A Kaleidoscope of Keratin Gene Expression and the Mosaic of Its Regulatory Mechanisms. Int J Mol Sci. Mar 15 2023;24(6)doi:10.3390/ijms24065603

69. Hemmaphan S, Bordeerat NK. Genotoxic Effects of Lead and Their Impact on the Expression of DNA Repair Genes. Int J Environ Res Public Health. Apr 3 2022;19(7)doi:10.3390/ijerph19074307

70. Meng L, Person RE, Beaudet AL. Ube3a-ATS is an atypical RNA polymerase II transcript that represses the paternal expression of Ube3a. Human molecular genetics. Jul 1 2012;21(13):3001–12. doi:10.1093/hmg/dds130

71. Hanna CW, Kelsey G. Features and mechanisms of canonical and noncanonical genomic imprinting. Genes & development. Jun 2021;35(11-12):821–834. doi:10.1101/gad.348422.121

72. Watanabe T, Tomizawa S, Mitsuya K, et al. Role for piRNAs and noncoding RNA in de novo DNA methylation of the imprinted mouse Rasgrf1 locus. Science. May 2011;332(6031):848–52. doi:10.1126/science.1203919

73. Perera BPU, Tsai ZT, Colwell ML, et al. Somatic expression of piRNA and associated machinery in the mouse identifies short, tissue-specific piRNA. Epigenetics. Apr 8 2019:1–18. doi:10.1080/15592294.2019.1600389

74. Fernandez-Medarde A, Santos E. The RasGrf family of mammalian guanine nucleotide exchange factors. Biochim Biophys Acta. Apr 2011;1815(2):170–88. doi:10.1016/j.bbcan.2010.11.001

75. McCarthy MM, Arnold AP. Reframing sexual differentiation of the brain. Nat Neurosci. Jun 2011;14(6):677–83. doi:10.1038/nn.2834

76. Bakulski KM, Rozek LS, Dolinoy DC, Paulson HL, Hu H. Alzheimer’s disease and environmental exposure to lead: the epidemiologic evidence and potential role of epigenetics. Curr Alzheimer Res. Jun 2012;9(5):563–73. doi:10.2174/156720512800617991

77. Banda N, Soe NC, Yabe J, et al. Sex dependent intergenerational effects of lead in mouse model. Sci Rep. Dec 4 2024;14(1):30233. doi:10.1038/s41598-024-81839-4

78. Wang K, Liu S, Svoboda LK, et al. Tissue– and Sex-Specific DNA Methylation Changes in Mice Perinatally Exposed to Lead (Pb). Front Genet. 2020;11:840. doi:10.3389/fgene.2020.00840

