## Supplementary Information for "Perinatal Lead (Pb) Exposure Increases Mouse Embryonic Weight and Alters Neuronal Gene Expression"

**Supporting Information**

Bambarendage P. U. Perera*[,1,2], Minghua Li[3], Anagha Tapaswi[1], Junru Pan[4], Dongyue Wang[1], Tejas Goswami[5], Rachel K. Morgan[1], Kelly M. Bakulski[6], Jaclyn M. Goodrich[1], Maureen A. Sartor[3,7], Dana C. Dolinoy[1,4], Justin A. Colacino*[,1,4]

[1]Department of Environmental Health Sciences, School of Public Health, University of Michigan, Ann Arbor, MI 48109, USA

[2]Division of Environmental Health Sciences, School of Public Health, University of California, Berkeley, Berkeley, CA 94704, USA

[3]Department of Computational Medicine and Bioinformatics, Medical School, University of Michigan, Ann Arbor, MI 48109, USA

[4]Department of Nutritional Sciences, School of Public Health, University of Michigan, Ann Arbor, MI 48109, USA

[5]Department of Chemistry, College of Literature, Science, and the Arts, University of Michigan, Ann Arbor, MI 48109, USA

[6]Department of Epidemiology, School of Public Health, University of Michigan, Ann Arbor, MI 48109, USA

[7]Department of Biostatistics, School of Public Health, University of Michigan, Ann Arbor, MI 48109, USA

**Table of contents**

Supplementary Figures S1–S16

Supplementary Table Information

### SUPPLEMENTARY FIGURES

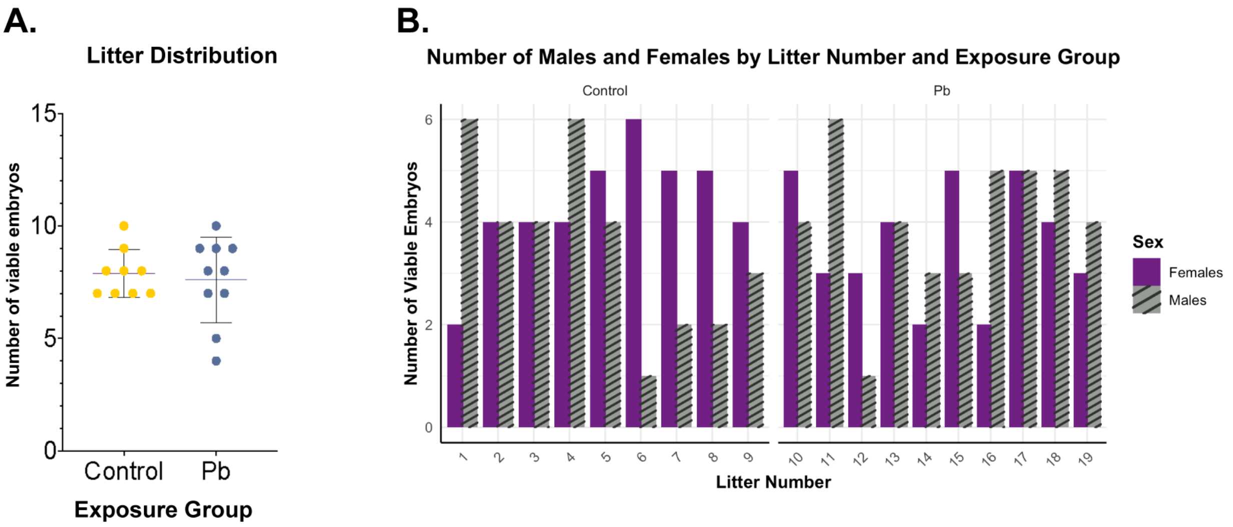


**Supplementary Figure S1**. **Litter and sex distribution for the perinatal Pb exposure study.** The figure shows results related to $n$=19 litters and $n$=147 total samples. (**A**) The litter distribution for control (yellow) and Pb (dark blue) exposure groups. The control group contained a total of 9 litters between 7–10 viable embryos per litter, while the Pb-exposed group contained a total of 10 litters at 4–10 embryos per litter. (**B**) The sex distribution for control (litter numbers 1–9) and Pb (litter numbers 10–19) exposure groups based on sex genotyping of the embryonic sac tissues. Viable embryos for females and males are represented by purple and grey striped bars, respectively.

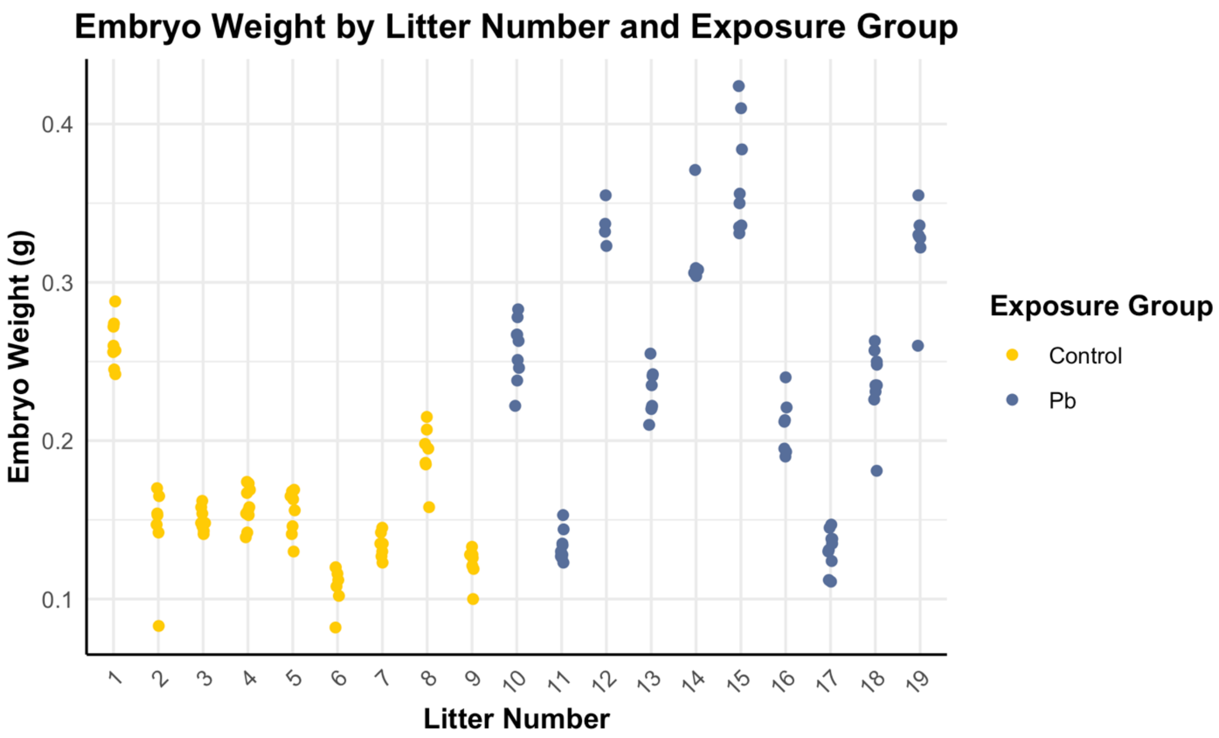


**Supplementary Figure S2**. **Whole embryo weight based on exposure group and litter.** The scatter plot represents raw data (*n*=146; excluding *n*=1 control embryo weight) used for the combined sex model indicating control (yellow; litter numbers 1–9) and Pb exposure (dark blue; litter numbers 10–19) groups separated by litter.

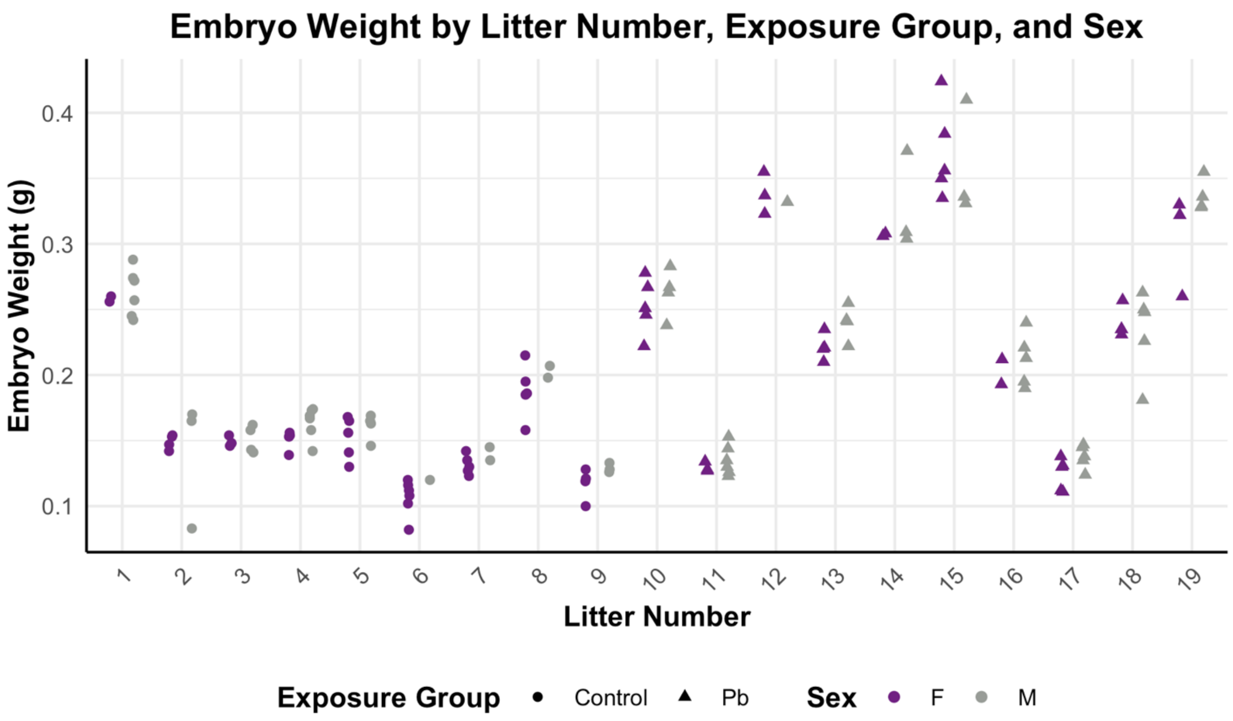


**Supplementary Figure S3**. **Whole embryo weight based on exposure group, sex, and litter.** The scatter plot represents raw data (*n*=146; excluding *n*=1 control male embryo weight) used for the sex stratified model indicating control (litter numbers 1–9) and Pb (litter numbers 10–19) exposure groups separated by litter for females (F; purple) and males (M; grey).

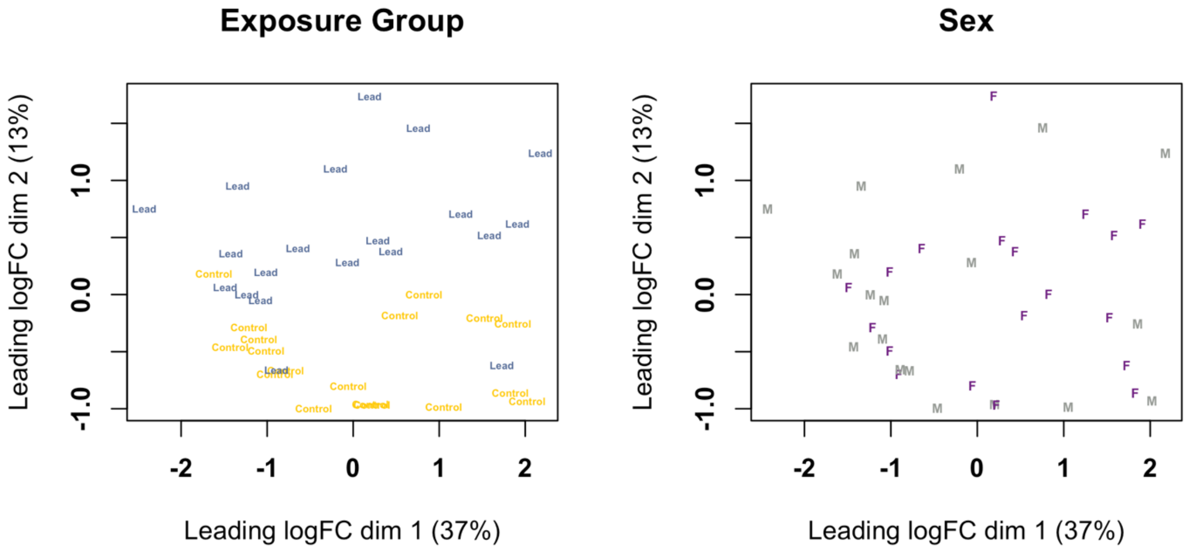


**Supplementary Figure S4**. **Sample variability visualized by MDS plots for all embryo head samples at E13–15.** The MDS plots represent information related to *n*=38 RNA-seq samples. The left panel indicates the sample distribution and variability based on control (yellow; *n*=18) and Pb (dark blue; *n*=20) exposure groups. The right panel indicates the sample distribution and variability based on sex (female: F; purple; *n*=19 and male: M; grey; *n*=19).

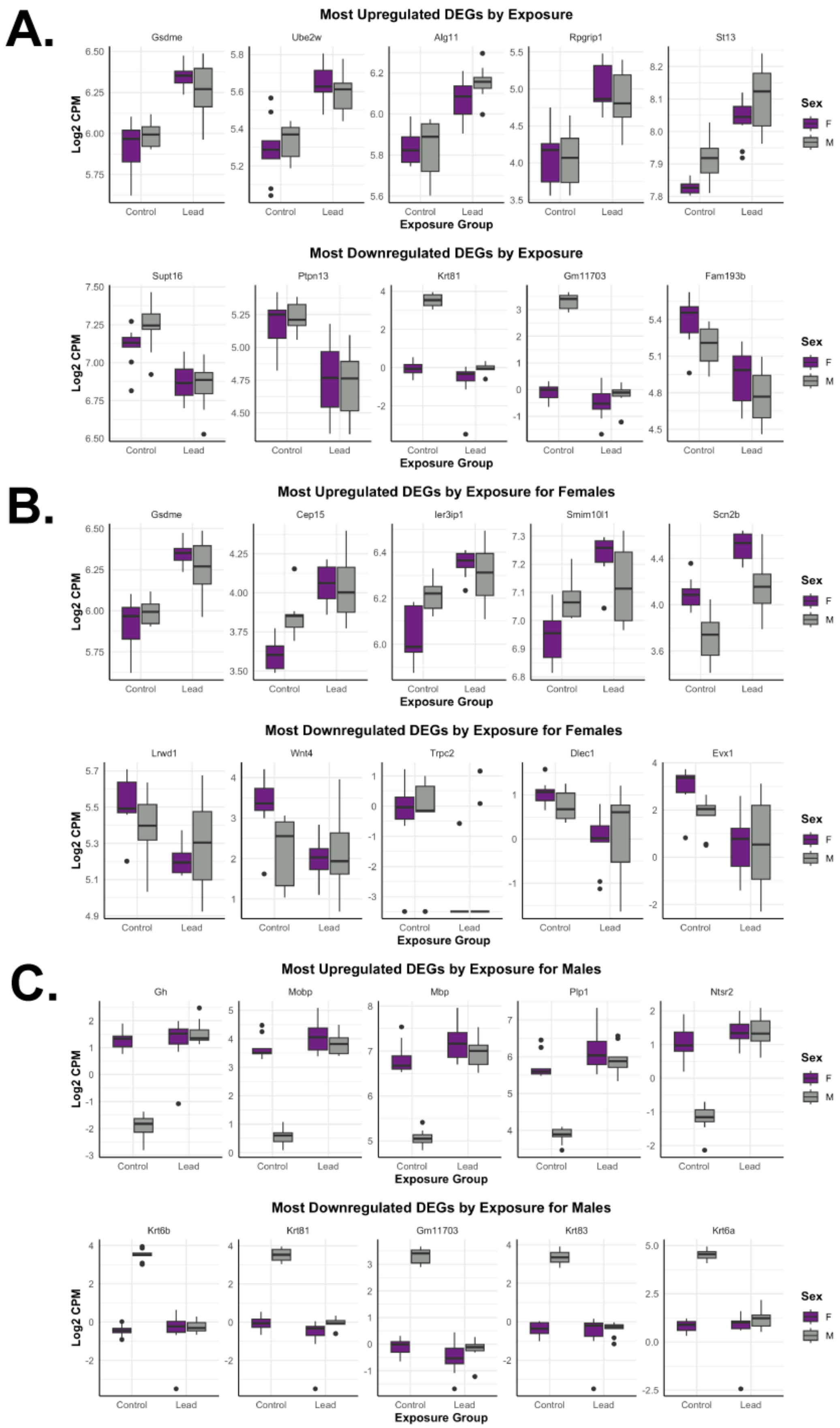
A.
Most Upregulated DEGs by Exposure
Gsdme
Ube2w
Alg11
Rpgrip1
St13
Log2 CPM
Control
Lead
Exposure Group
Sex
F
M
Most Downregulated DEGs by Exposure
Supt16
Ptpn13
Krt81
Gm11703
Fam193b
B.
Most Upregulated DEGs by Exposure for Females
Gsdme
Cep15
Ier3ip1
Smim10l1
Scn2b
Most Downregulated DEGs by Exposure for Females
Lrwd1
Wnt4
Trpc2
Dlec1
Evx1
C.
Most Upregulated DEGs by Exposure for Males
Gh
Mobp
Mbp
Plp1
Ntsr2
Most Downregulated DEGs by Exposure for Males
Krt6b
Krt81
Gm11703
Krt83
Krt6a

**Supplementary Figure S5**. **Differentially expressed genes based on exposure group and stratified by sex.** The combined model shows results related to *n*=38 RNA-seq samples, including *n*=18 for control and *n*=20 for the Pb exposure groups. The female/male models show results related to *n*=19 RNA-seq samples, including *n*=9 for control and *n*=10 for the Pb exposure groups. The boxplots represent gene expression data for females (F; purple) and males (M; grey). (**A**) The most upregulated *(Gsdme*, *Ube2w*, *Alg11*, *Rpgrip1*, and *St13*; upper panel) and downregulated (*Supt16*, *Ptpn13*, *Krt81*, *Gm11703*, and *Fam193b;* bottom panel) differentially expressed genes (based on FDR) by Pb exposure, stratified by sex, for the combined model. (**B**) The most upregulated (*Gsdme*, *Cep15*, *Ier3ip1*, *Smim10l1*, and *Scn2b*; upper panel) and downregulated (*Lrwd1*, *Wnt4*, *Trpc2*, *Dlec1*, and *Evx1*; bottom panel) differentially expressed genes (based on FDR) by Pb exposure, stratified by sex, for the female model. (**C**) The most upregulated (*Gh*, *Mobp*, *Mbp*, *Plp1*, and *Ntsr2*; upper panel) and downregulated (*Krt6b*, *Krt81*, *Gm11703*, *Krt83*, and *Krt6a*; bottom panel) differentially expressed genes (based on FDR) by Pb exposure, stratified by sex, for the male model. DEGs: differentially expressed genes.

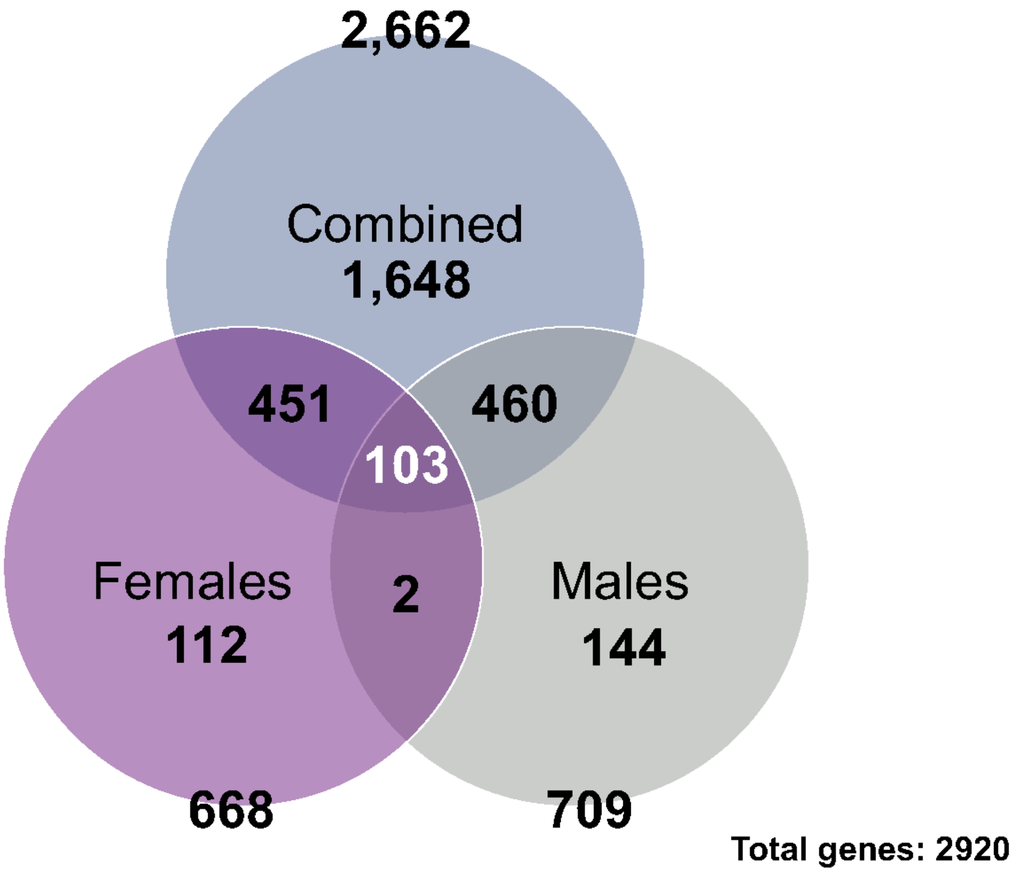


**Supplementary Figure S6. Combined sex, Female, and Male model-specific differentially expressed gene comparisons.** The combined model shows 2,662 differentially expressed genes related to *n*=38 RNA-seq samples, including *n*=18 for control and *n*=20 for the Pb exposure groups. The female/male models show 668/709 differentially expressed genes, respectively, related to *n*=19 RNA-seq samples, including *n*=9 for control and *n*=10 for the Pb exposure groups. Collectively, 2,920 differentially expressed genes were dysregulated by gestational exposure to Pb in the embryonic head.

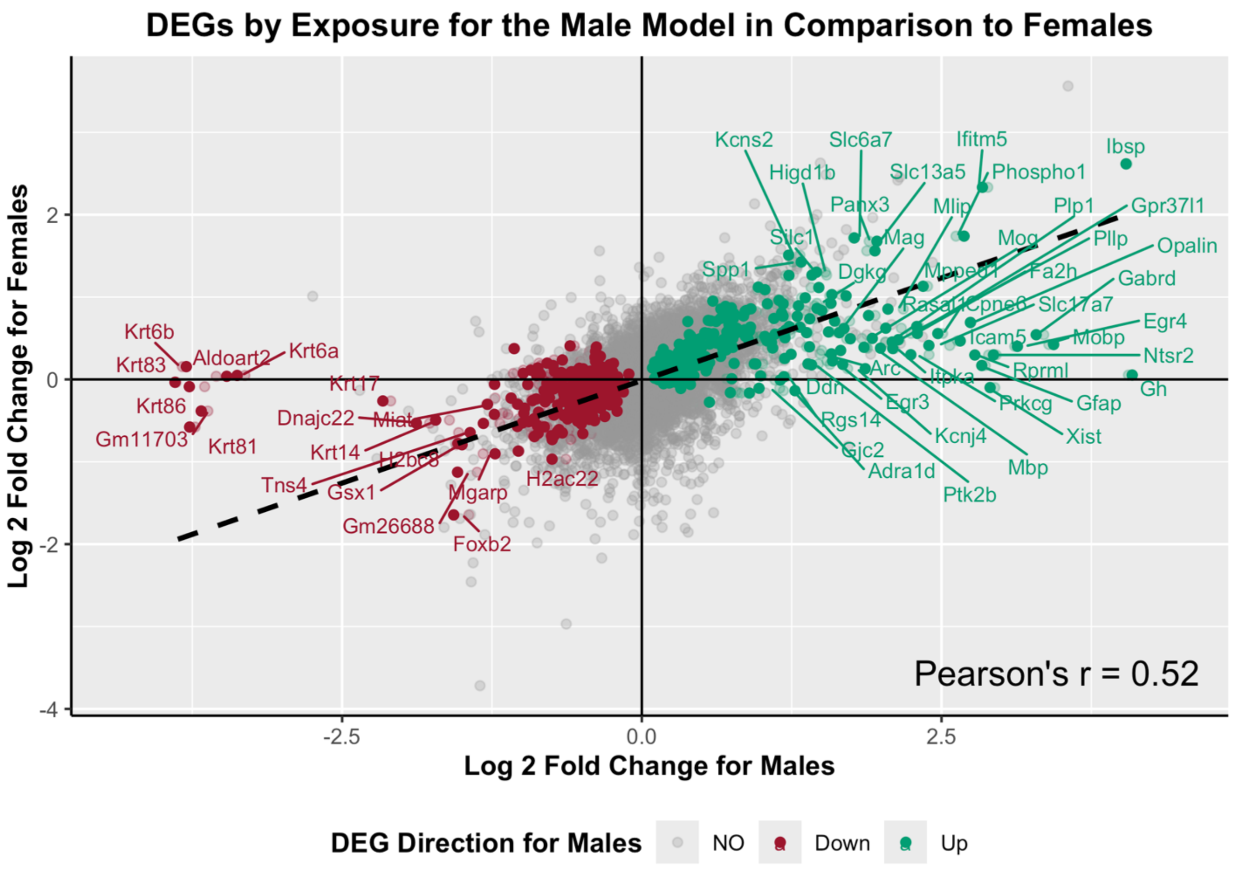


**Supplementary Figure S7**. **Correlation of differentially expressed genes between female vs. male models of Pb exposure.** The female model shows 668 differentially expressed genes, while the male models show 709 differentially expressed genes based on exposure to Pb, representing *n*=19 RNA-seq samples, including *n*=9 for control and *n*=10 for the Pb exposure groups for each sex. The graph shows the LogFC between female and male gene expression. The data indicates upregulated (teal; LogFC>0; FDR<0.05) and downregulated (red; LogFC<0; FDR<0.05) differentially expressed genes in Pb-exposed males compared to females, relative to their control counterparts. The trendline represents the Pearson's correlation coefficient calculation. DEGs: differentially expressed genes.

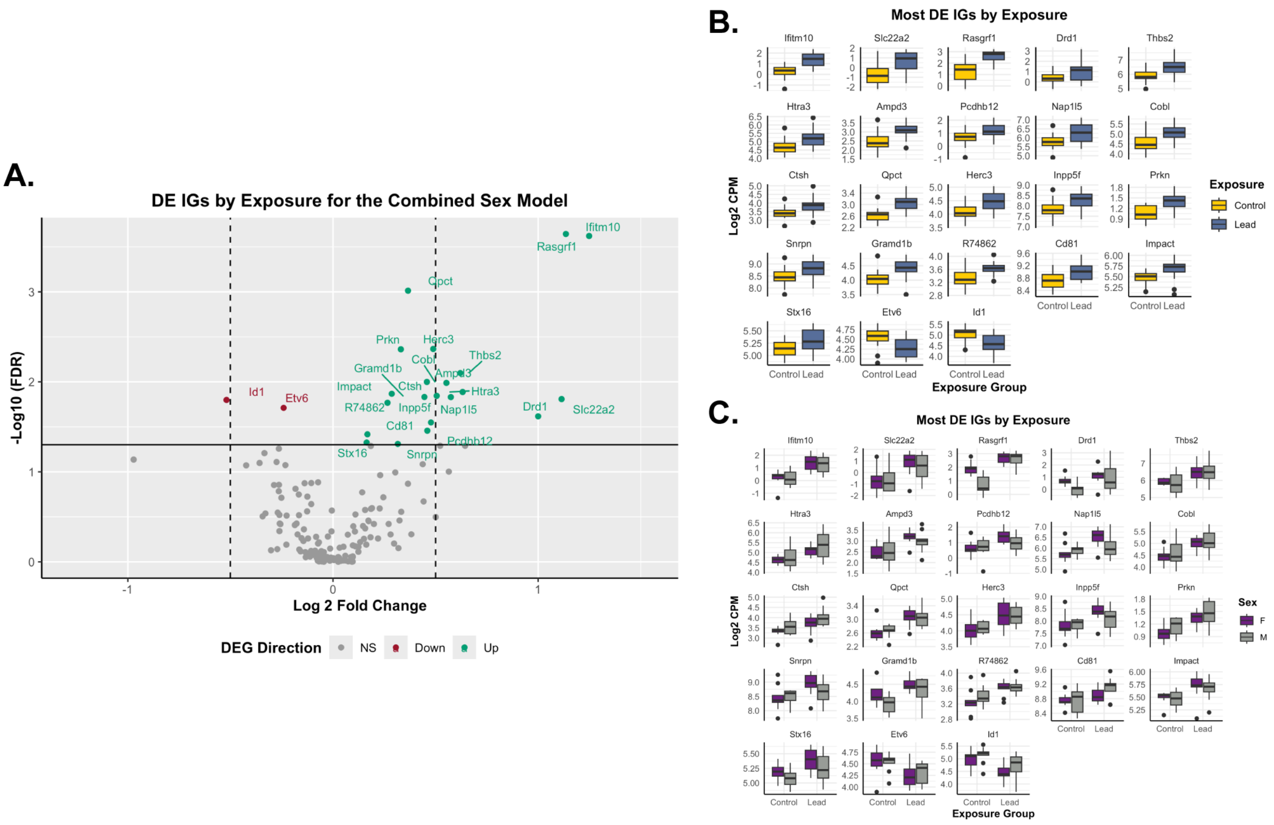


**Supplementary Figure S8**. **Differentially expressed imprinted genes based on exposure group for the combined sex model.** (**A**) The volcano plot represents upregulated (teal; LogFC>0; FDR<0.05) and downregulated (red; LogFC<0; FDR<0.05) differentially expressed imprinted genes by Pb exposure relative to control. The dotted lines indicate 0.5 LogFC, while non-significant (NS) genes are shown in grey. (**B**) The differentially expressed imprinted genes (*Ifitm10*, *Slc22a2*, *Rasgrf1*, *Drd1*, *Thbs2*, *Htra3*, *Ampd3*, *Pcdhb12*, *Nap1l5*, *Cobl*, *Ctsh*, *Qpct*, *Herc3*, *Inpp5f*, *Prkn*, *Snrpn*, *Gramd1b*, *R74862*, *Cd81*, *Impact*, *Stx16*, *Etv6*, and *Id1*; organized based on LogFC in descending order; FDR<0.05) based on exposure group (control: yellow; and Pb: dark blue). (**C**) Differentially expressed imprinted genes by Pb exposure, stratified by sex, for the combined model. The boxplots represent gene expression results for all tested animals including females (F; purple) and males (M; grey) for the imprinted genes listed above. DE IGs: differentially expressed imprinted genes.

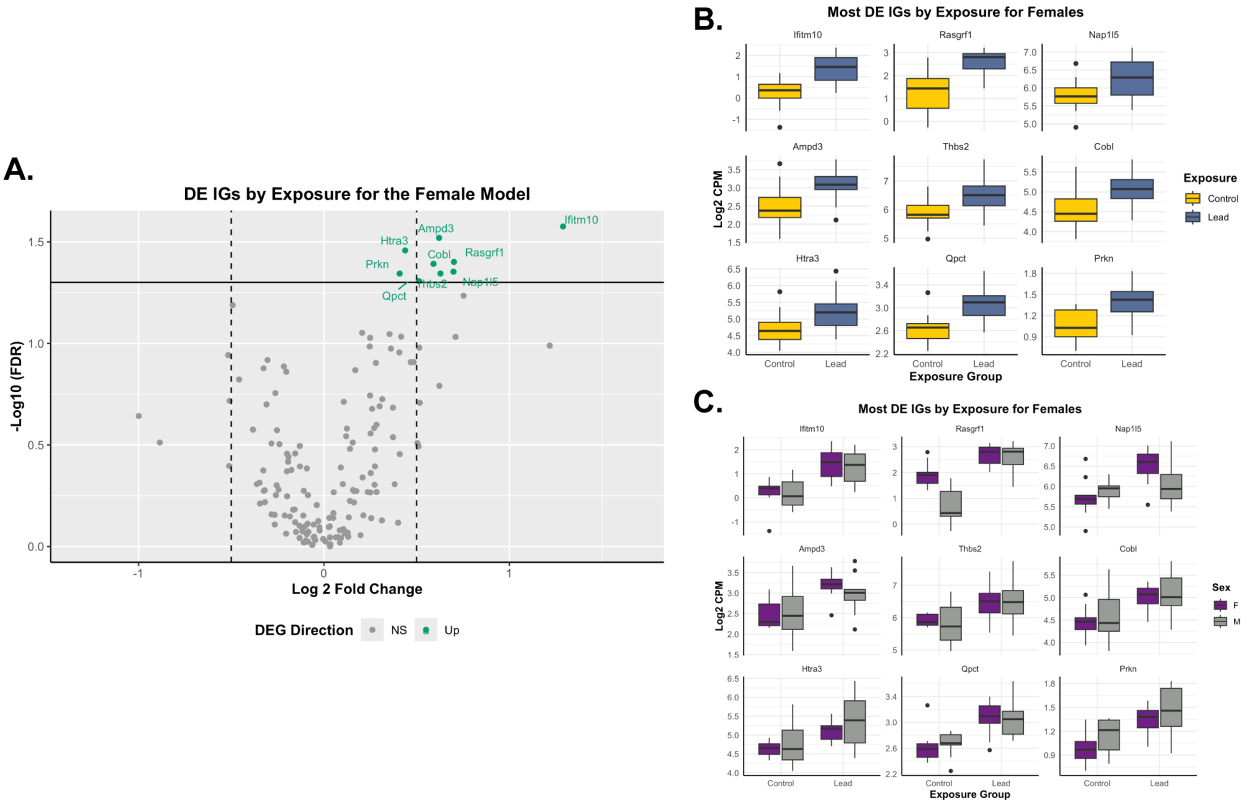


**Supplementary Figure S9**. **Differentially expressed imprinted genes based on exposure group for the female model.** (**A**) The volcano plot represents upregulated (teal; LogFC>0; FDR<0.05) differentially expressed imprinted genes by Pb-exposed females relative to control females. The dotted lines indicate 0.5 LogFC, while non-significant (NS) genes are shown in grey. (**B**) The differentially expressed imprinted genes (*Ifitm10*, *Rasgrf1*, *Nap1l5*, *Ampd3*, *Thbs2*, *Cobl*, *Htra3*, *Qpct*, and *Prkn*; organized based on LogFC in descending order; FDR<0.05) based on exposure group (control: yellow; and Pb: dark blue) for females. (**C**) Differentially expressed imprinted genes by Pb exposure, stratified by sex for females, are compared to the combined model to visualize sex-specific differences. The boxplots represent gene expression results for females (F; purple) and males (M; grey) for the imprinted genes listed above. DE IGs: differentially expressed imprinted genes.

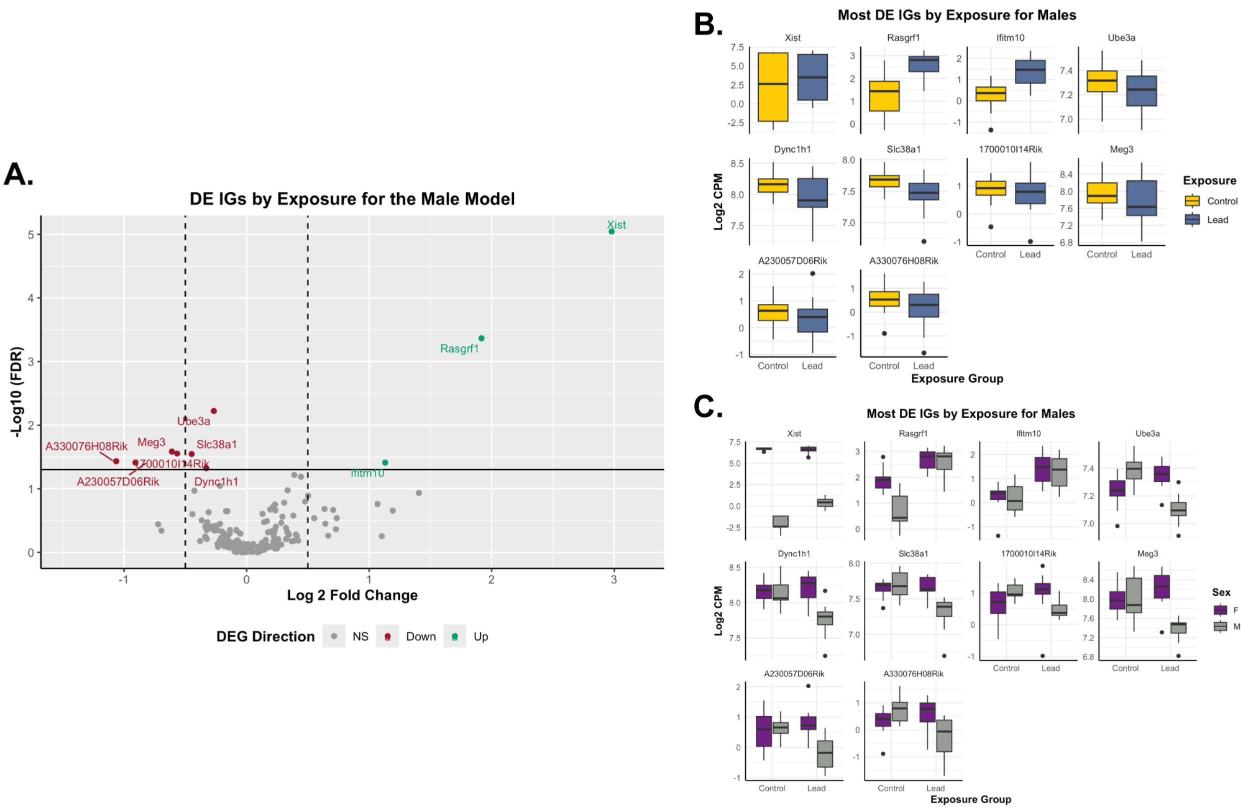


**Supplementary Figure S10**. **Differentially expressed imprinted genes based on exposure group for the male model.** (**A**) The volcano plot represents upregulated (teal; LogFC>0; FDR<0.05) and downregulated (red; LogFC<0; FDR<0.05) differentially expressed imprinted genes by Pb-exposed males, relative to control males. The dotted lines indicate 0.5 LogFC, while non-significant (NS) genes are shown in grey. (**B**) The differentially expressed imprinted genes (*Xist*, *Rasgrf1*, *Ifitm10*, *Ube3a*, *Dync1h1*, *Slc38a1*, *1700010I14Rik*, *Meg3*, *A230057D06Rik*, and *A330076H08Rik*; organized based on LogFC in descending order; FDR<0.05) based on exposure group (control: yellow; and Pb: dark blue) for males. (**C**) Differentially expressed imprinted genes by Pb exposure for males, stratified by sex, are compared to the combined model to visualize sex-specific differences. The boxplots represent gene expression results for females (F; purple) and males (M; grey) for the imprinted genes listed above. DE IGs: differentially expressed imprinted genes.

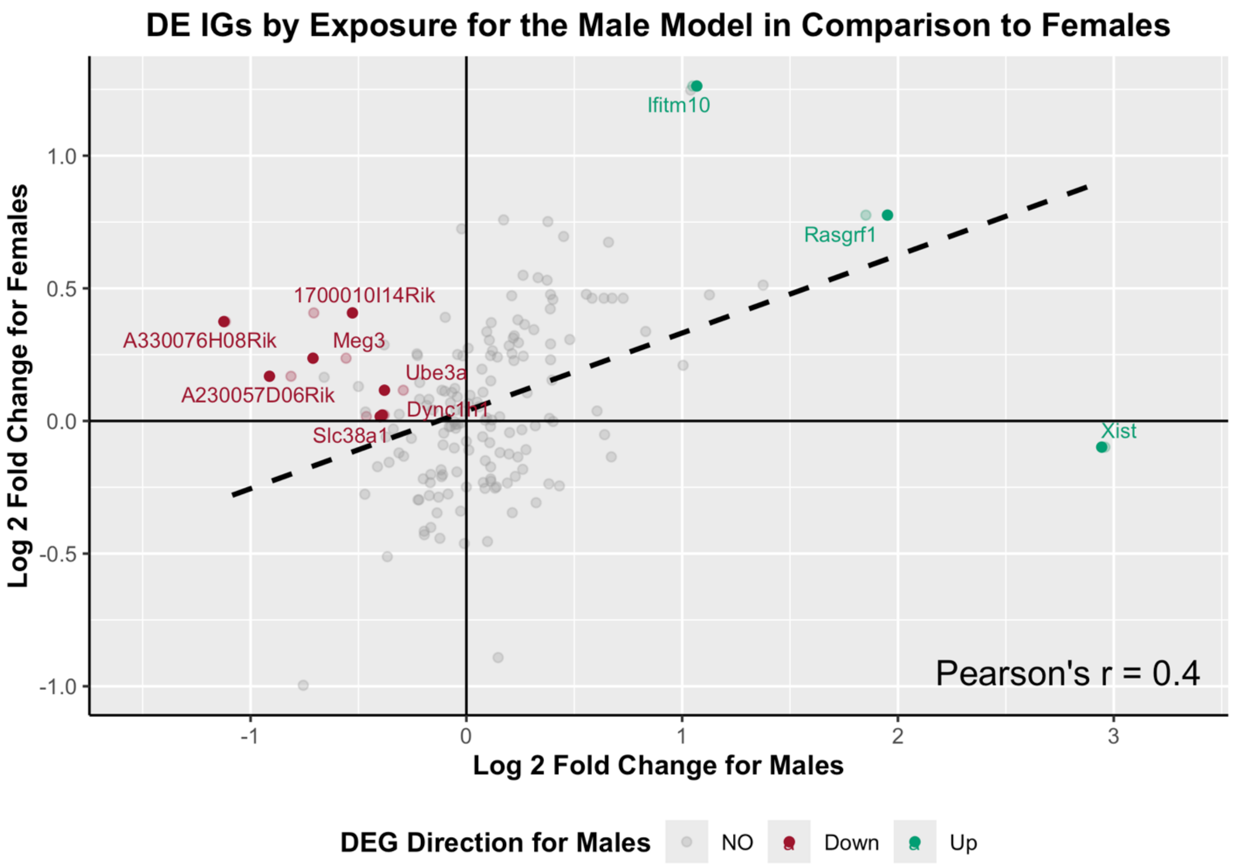


**Supplementary Figure S11**. **Correlation of differentially expressed imprinted genes between female vs. male models of Pb exposure.** The female model shows 9 differentially expressed imprinted genes, while the male models show 10 differentially expressed imprinted genes based on exposure to Pb, representing *n*=19 RNA-seq samples, including *n*=9 for control and *n*=10 for the Pb exposure groups for each sex. The graph shows the LogFC between female and male imprinted gene expression. The data indicates upregulated (teal; LogFC>0; FDR<0.05) and downregulated (red; LogFC<0; FDR<0.05) differentially expressed imprinted genes in Pb-exposed males compared to females, relative to their control counterparts. The trendline represents the Pearson's correlation coefficient calculation. DE IGs: differentially expressed imprinted genes.

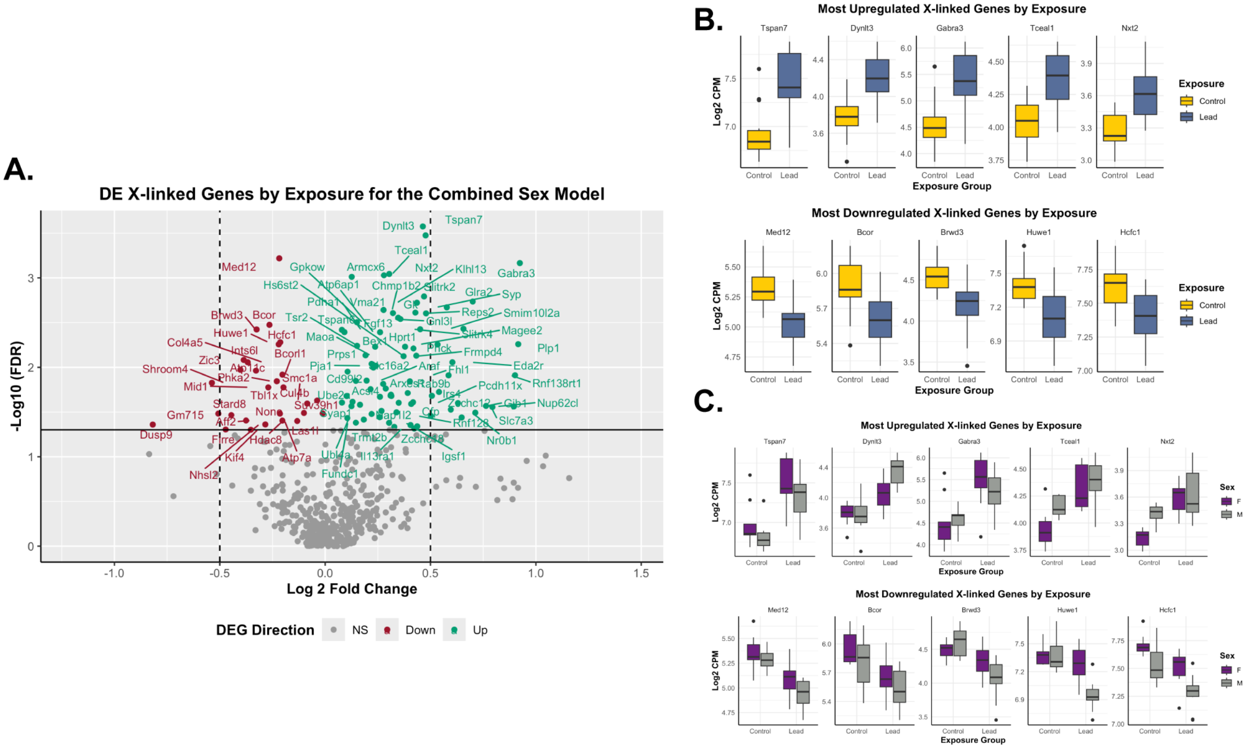


**Supplementary Figure S12**. **Differentially expressed X-linked genes based on exposure group for the combined sex model.** (**A**) The volcano plot represents upregulated (teal; LogFC>0; FDR<0.05) and downregulated (red; LogFC<0; FDR<0.05) differentially expressed X-linked genes by Pb exposure relative to control. The dotted lines indicate 0.5 LogFC, while non-significant (NS) genes are shown in grey. (**B**) The most upregulated (*Tspan7*, *Dynlt3*, *Gabra3*, *Tceal1*, and *Nxt2*; upper panel) and downregulated (*Med12*, *Bcor*, *Brwd3*, *Huwe1*, and *Hcfc1*; bottom panel) differentially expressed X-linked genes (based on FDR) by exposure group (control: yellow; and Pb: dark blue). (**C**) The most up (upper panel) and downregulated (bottom panel) differentially expressed X-linked genes by Pb exposure, stratified by sex, for the combined model. The boxplots represent gene expression results for all tested animals including females (F; purple) and males (M; grey) for the X-linked genes listed above. DE: differentially expressed.

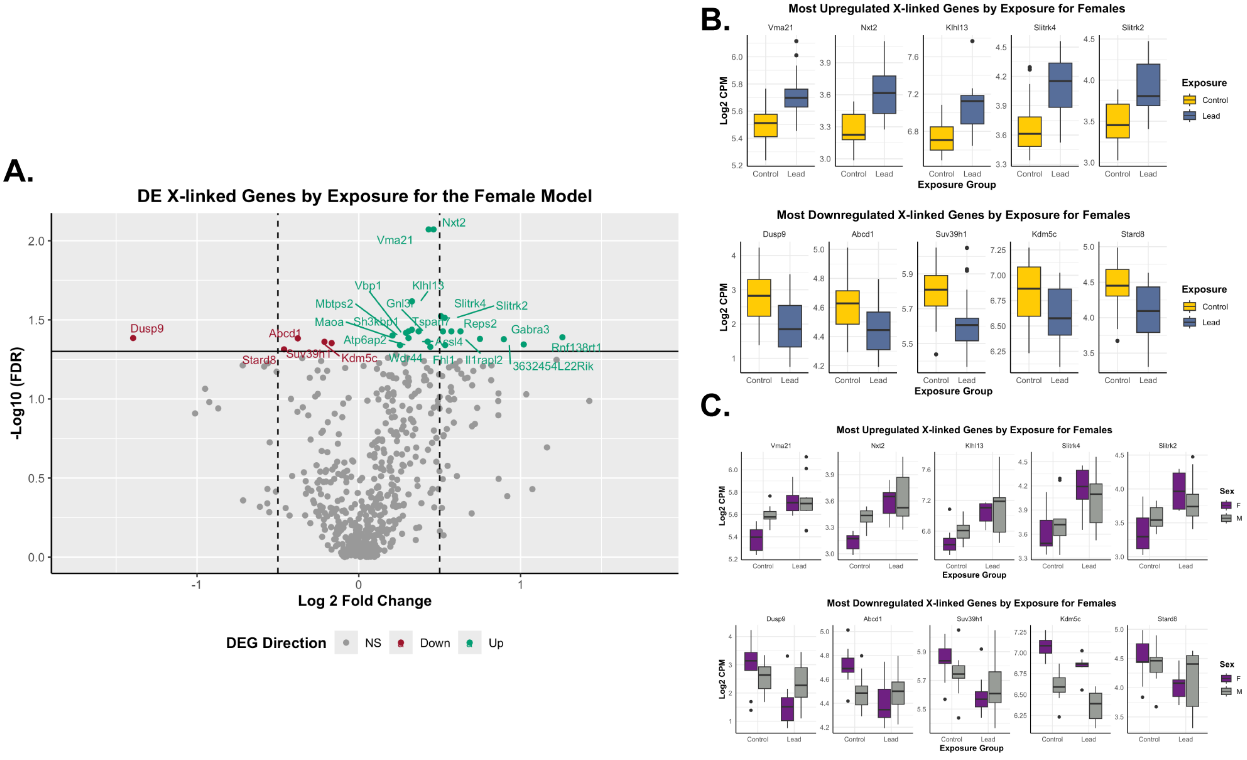


**Supplementary Figure S13. Differentially expressed X-linked genes based on exposure group for the female model.** (**A**) The volcano plot represents upregulated (teal; LogFC>0; FDR<0.05) and downregulated (red; LogFC<0; FDR<0.05) differentially expressed X-linked genes by Pb-exposed females, relative to control females. The dotted lines indicate 0.5 LogFC, while non-significant (NS) genes are shown in grey. (**B**) The most upregulated (*Vma21*, *Nxt2*, *Klhl13*, *Slitrk4*, and *Slitrk2*; upper panel) and downregulated (*Dusp9*, *Abcd1*, *Suv39h1*, *Kdm5c*, and *Stard8*; bottom panel) differentially expressed X-linked genes (based on FDR) by exposure group (control: yellow; and Pb: dark blue) for females. (**C**) The most up (upper panel) and downregulated (bottom panel) differentially expressed X-linked genes by Pb exposure for females, stratified by sex, are compared to the combined model to visualize sex-specific differences. The boxplots represent gene expression results for all tested animals including females (F; purple) and males (M; grey) for the X-linked genes listed above. DE: differentially expressed.

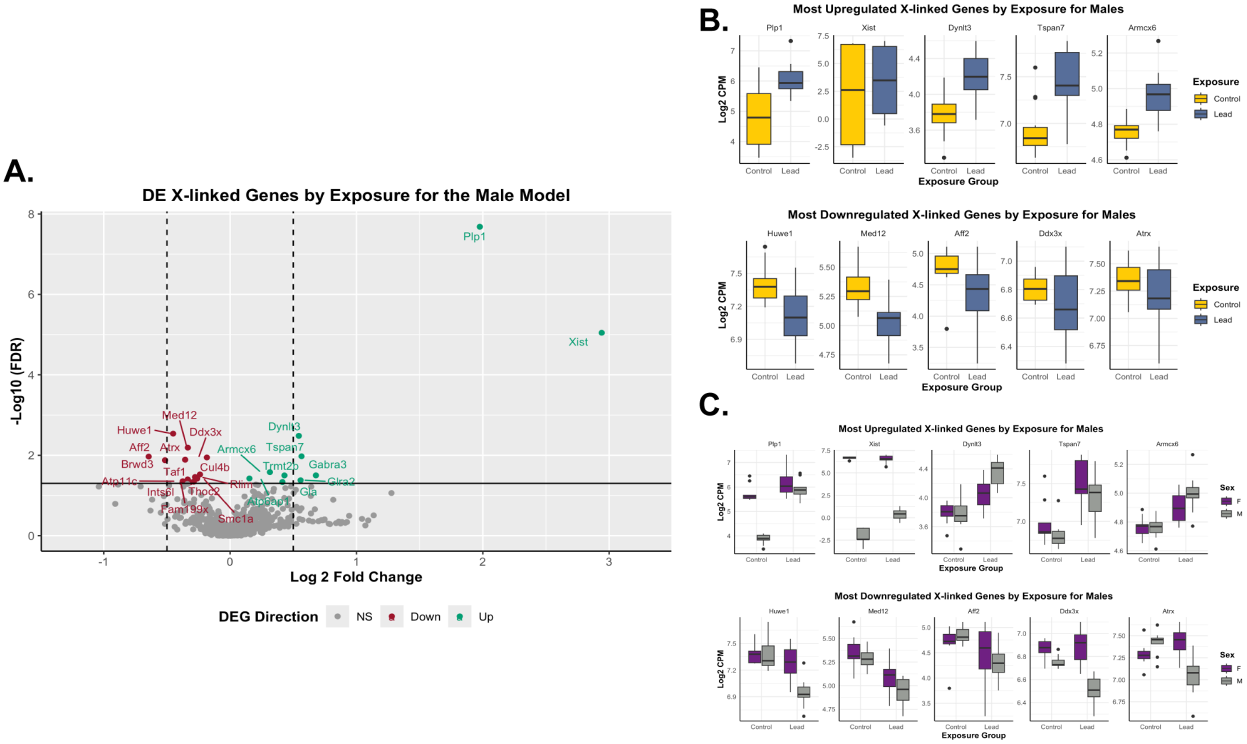


**Supplementary Figure S14**. **Differentially expressed X-linked genes based on exposure group for the male model.** (**A**) The volcano plot represents upregulated (teal; LogFC>0; FDR<0.05) and downregulated (red; LogFC<0; FDR<0.05) differentially expressed X-linked genes by Pb-exposed males, relative to control males. The dotted lines indicate 0.5 LogFC, while non-significant (NS) genes are shown in grey. (**B**) The most upregulated (*Plp1*, *Xist*, *Dynlt3*, *Tspan7*, and *Armcx6*; upper panel) and downregulated (*Huwe1*, *Med12*, *Aff2*, *Ddx3x*, and *Atrx*; bottom panel) differentially expressed X-linked genes (based on FDR) by exposure group (control: yellow; and Pb: dark blue) for males. (**C**) The most up (upper panel) and downregulated (bottom panel) differentially expressed X-linked genes by Pb exposure for males, stratified by sex, are compared to the combined model to visualize sex-specific differences. The boxplots represent gene expression results for all tested animals including females (F; purple) and males (M; grey) for the X-linked genes listed above. DE: differentially expressed.

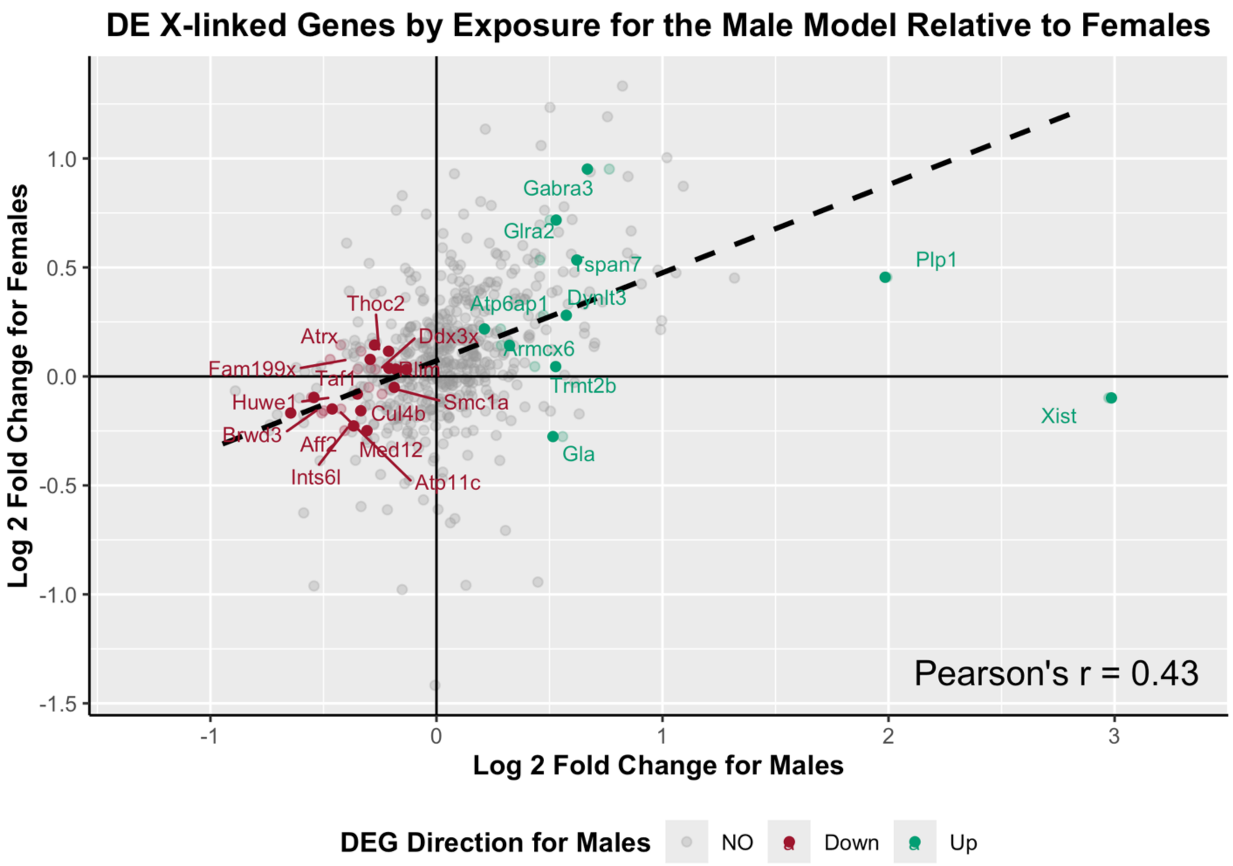


**Supplementary Figure S15**. **Correlation of differentially expressed X-linked genes between female vs. male models of Pb exposure.** The female model shows 27 differentially expressed X-linked genes, while the male models show 24 differentially expressed X-linked genes based on exposure to Pb, representing $n$=19 RNA-seq samples, including $n$=9 for control and $n$=10 for the Pb exposure groups for each sex. The graph shows the LogFC between females and males X-linked gene expression. The data indicates upregulated (teal; LogFC>0; FDR<0.05) and downregulated (red; LogFC<0; FDR<0.05) differentially expressed X-linked genes in Pb-exposed males compared to females, relative to their control counterparts. The trendline represents the Pearson's correlation coefficient calculation. DE: differentially expressed.

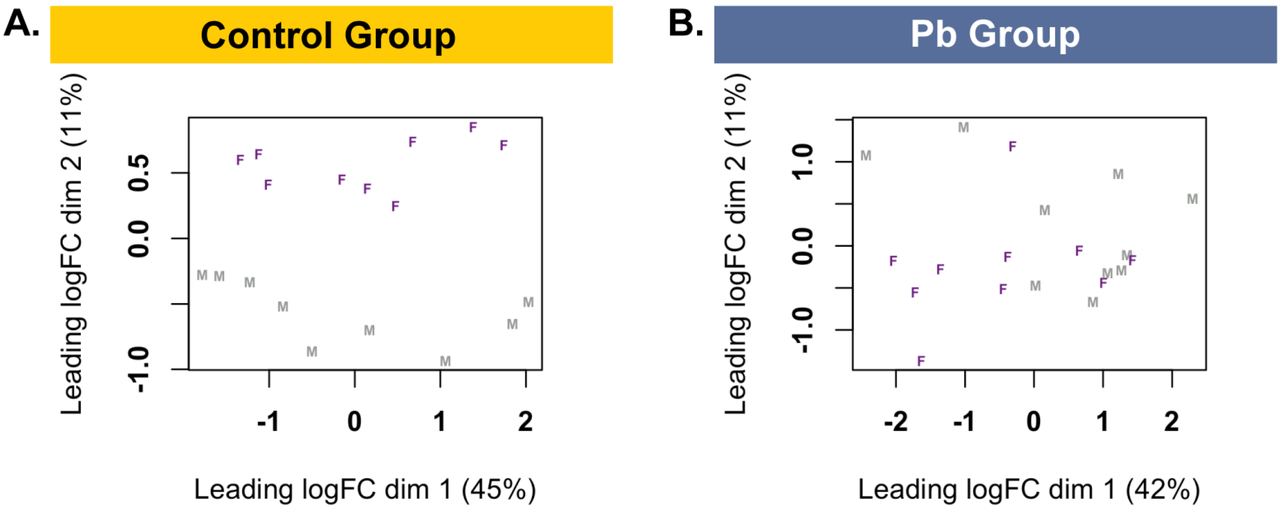


**Supplementary Figure S16**. **Sex-specific sample variability visualized by MDS plots for each exposure group.** (**A**) The MDS plots represent information related to *n*=9 females (F; purple) and *n*=9 males (M; grey) RNA-seq samples. The sample distribution and variability are specifically assessed for the control group (yellow; *n*=18). (**B**) The MDS plots show results related to *n*=10 females (F; purple) and *n*=10 males (M; grey) RNA-seq samples from the Pb group. The sample distribution and variability are specifically assessed for the Pb (dark blue; *n*=20) exposure group.

### SUPPLEMENTARY TABLE INFORMATION

**Table S1**. RNA-seq Library Sizes and Metadata for the Embryo Head at E13–15.

**Table S2**. Number of Differentially Expressed Genes with |LogFC|>0 and FDR<0.05 for Combined Sex.

**Table S3**. Number of Differentially Expressed Genes with |LogFC|>0 and FDR<0.05 for Females.

**Table S4**. Number of Differentially Expressed Genes with |LogFC|>0 and FDR<0.05 for Males.

**Table S5**. Gene Set Enrichment Analysis by LRpath for Combined Sex.

**Table S6**. KEGG Analysis by LRpath for Combined Sex.

**Table S7**. Gene Set Enrichment Analysis by LRpath for Females.

**Table S8**. KEGG Analysis by LRpath for Females.

**Table S9**. Gene Set Enrichment Analysis by LRpath for Males.

**Table S10**. KEGG Analysis by LRpath for Males.

**Table S11**. Number of Imprinted Genes Among Differentially Expressed Genes with |LogFC|>0 and FDR<0.05 for Combined Sex.

**Table S12**. Number of Imprinted Genes Among Differentially Expressed Genes with |LogFC|>0 and FDR<0.05 for Females.

**Table S13**. Number of Imprinted Genes Among Differentially Expressed Genes with |LogFC|>0 and FDR<0.05 for Males.

**Table S14**. Number of X-linked Genes Among Differentially Expressed Genes with |LogFC|>0 and FDR<0.05 for Combined Sex.

**Table S15**. Number of X-linked Genes Among Differentially Expressed Genes with |LogFC|>0 and FDR<0.05 for Females.

**Table S16**. Number of X-linked Genes Among Differentially Expressed Genes with |LogFC|>0 and FDR<0.05 for Males.

**Table S17**. Number Differentially Expressed Genes with |LogFC|>0 and FDR<0.05 for the Control Group (Male Relative to Female).
